# Mosaic H3K9me3 at BREACHes predicts synaptic gene expression associated with fragile X syndrome cognitive severity

**DOI:** 10.1101/2025.03.19.644148

**Authors:** Kenneth Pham, Thomas Malachowski, Linda Zhou, Ji Hun Kim, Chuanbin Su, Jennifer E. Phillips-Cremins

**Affiliations:** Department of Genetics, Perelman School of Medicine, University of Pennsylvania, Philadelphia, PA; Department of Bioengineering, School of Engineering and Applied Sciences, University of Pennsylvania, Philadelphia, PA, USA; Epigenetics Institute, Perelman School of Medicine, University of Pennsylvania, Philadelphia, PA, USA

## Abstract

Diseases vary in clinical presentation across individuals despite the same molecular diagnosis. In fragile X syndrome (FXS), mutation-length expansion of a CGG short tandem repeat (STR) in *FMR1* causes reduced gene expression and FMRP loss. Nevertheless, *FMR1* and FMRP are limited predictors of adaptive functioning and cognition in FXS patients, suggesting that molecular correlates of clinical measures would add diagnostic value. We recently uncovered Megabase-scale domains of heterochromatin (BREACHes) in FXS patient-derived iPSCs. Here, we identify BREACHes in FXS brain tissue (N=4) and absent from sex/age-matched neurotypical controls (N=4). BREACHes span >250 genes and exhibit patient-specific H3K9me3 variation. Using N=4 FXS iPSC lines and N=7 single-cell isogenic FXS iPSC subclones, we observe a strong correlation between inter-sample H3K9me3 variation and heterogeneous BREACH gene repression. We demonstrate improved prediction of cognitive metrics in FXS patients with an additive model of blood FMRP and mRNA levels of H3K9me3-mosaic, but not H3K9me3-invariant, BREACH genes. Our results highlight the utility of H3K9me3 variation at BREACHes for identifying genes associated with FXS clinical metrics.

## Introduction

Fragile X syndrome (FXS) is the most common genetic form of inherited intellectual disability (*1*). FXS patients present at a young age with a range of neurological phenotypes, including developmental delay, seizures, anxiety, and movement disorders, as well as hallmark physical features, such as long face, prominent forehead and jaw, macroorchidism, connective tissue dysplasia, and joint hypermobility (*1–3*). The genetic cause of FXS is the mutation-length expansion of a CGG short tandem repeat (STR) tract in the 5’ untranslated region (5’UTR) of the *FMR1* gene (*4–8*). STR tract expansion beyond 200 CGG repeat units, termed full mutation-length, correlates with DNA methylation at the *FMR1* promoter, reduction in *FMR1* mRNA levels, and loss of the *FMR1* protein product termed fragile X messenger ribonucleoprotein (FMRP) (*9–13*). Thus, the classic model of FXS includes CGG expansion, local DNA methylation, and *FMR1* silencing.

DNA methylation and gene expression mosaicism have been reported across single cells (*14*) and patients with FXS (*15–22*). Case studies document FXS patients harboring alleles with an unmethylated *FMR1* promoter despite a full mutation-length CGG STR (*15–17*). Moreover, mosaicism in the methylation of the *FMR1* promoter is associated with less severe FXS clinical presentations (*18, 19*). Unmethylated *FMR1* promoters in FXS patients and the resultant higher levels of FMRP predict less severe symptoms (*15, 20–22*). Together, these data highlight the phenomena of epigenetic mosaicism and its link to the severity of FXS clinical presentations.

Genome-wide chromatin changes have recently been reported in FXS patient-derived cell lines (*23, 24*). Long-range looping interactions between *FMR1* and distal enhancers, as well as the synaptic genes *SLITRK2* and *SLITRK4,* are abolished in FXS patient-derived fibroblasts, induced pluripotent stem cells (iPSCs), iPSC-derived neural progenitor cells (iPSC-NPCs), and EBV-transformed B lymphoblasts (*23, 24*). *FMR1* loop disruption correlates with reduction in its mRNA levels. More recently, we reported Megabase (Mb) scale domains of the heterochromatic histone modification H3 lysine-9 tri-methylation (H3K9me3) on autosomes and the X-chromosome in FXS patient-derived fibroblasts, iPSCs, iPSC-NPCs, and EBV-transformed B lymphoblasts (*24*). Such domains – termed BREACHes (**<u>B</u>**eacons of **<u>R</u>**epeat **<u>E</u>**xpansion **<u>A</u>**nchored by **<u>C</u>**ontacting **<u>H</u>**eterochromatin) – correlate with the silencing of genes linked to cellular and tissue functions associated with FXS, such as synaptic plasticity, neural cell adhesion, reproductive development, and epithelial integrity (*24*). These data suggest that the link between heterochromatin and gene expression silencing in FXS might extend more broadly to other genes beyond only *FMR1*.

The extent to which H3K9me3 signal at BREACHes varies across FXS patients in tissue and the clinical relevance of heterochromatin mosaicism remains unknown. Here, we map H3K9me3 signal genome-wide in rare FXS and neurotypical brain tissue samples acquired from the NIH NeuroBioBank, identify FXS-specific BREACHes, and analyze H3K9me3 mosaicism across FXS patients. We identify BREACHes in male FXS patient brain samples (N=4) and devoid of H3K9me3 signal in age-/sex-matched neurotypical donors (N=4). BREACHes span >250 genes and exhibit patient-specific H3K9me3 variation. Using RNA-seq and H3K9me3 ChIP-seq data in NPCs derived from N=4 independent FXS iPSC lines (iPSC-NPCs) and N=7 single-cell isogenic subclones from a single FXS iPSC line (*24*), we observe a correlation between mosaic inter-sample H3K9me3 signal and heterogeneous mRNA levels across lines and clones. Using an additive model combining blood FMRP and blood mRNA levels, we discovered that H3K9me3-mosaic, but not H3K9me3-invariant, BREACH genes in FXS brain and iPSC-NPCs improved prediction of cognitive and adaptive functioning metrics in FXS patients compared to the current standard of FMRP levels alone. Together, our data suggest that H3K9me3 mosaicism at BREACHes can identify and prioritize synaptic genes with patient-to-patient mosaic repression that have predictive value over cognitive metrics clinically relevant for FXS diagnosis.

## Results

### Genome-wide identification of H3K9me3 BREACHes in FXS brain tissue

We previously reported Megabase-scale domains of H3K9me3 termed BREACHes in iPSC-NPCs from FXS patients and absent in sex-matched normal-length neurotypical individuals (*24*). In a pilot study, we confirmed that BREACHes exist in FXS brain tissue as well as cell lines, but a detailed characterization of BREACHes in tissue samples has not yet been conducted (*24*).

To identify H3K9me3-positive BREACHes across FXS patients, we performed H3K9me3 CUT&RUN in brain tissue samples from N=4 male FXS patients and N=4 sex- and age-matched neurotypical donors with no FXS diagnosis, including two previously published donor brain samples which we had not yet analyzed (**Table S1**). We identified H3K9me3 BREACHes present in at least 1 of 4 FXS and absent in 4 of 4 neurotypical brain tissue samples (**Fig. 1A**, **Table S2, Methods**). We observed that 3% of BREACHes (N=1/35) were unique to an individual FXS patient sample, 14% (N=13/35) were found in two FXS patient samples, about 75% (N=26/35) found in three samples, and about 8% (N=3/35) found in four out of four FXS patient brain samples (**Fig. 1B**). Our analyses identify BREACHes genome-wide unique to FXS brain tissue and suggest heterogeneity of H3K9me3 signal at BREACHes across FXS patient brain samples.

**Fig. 1.**
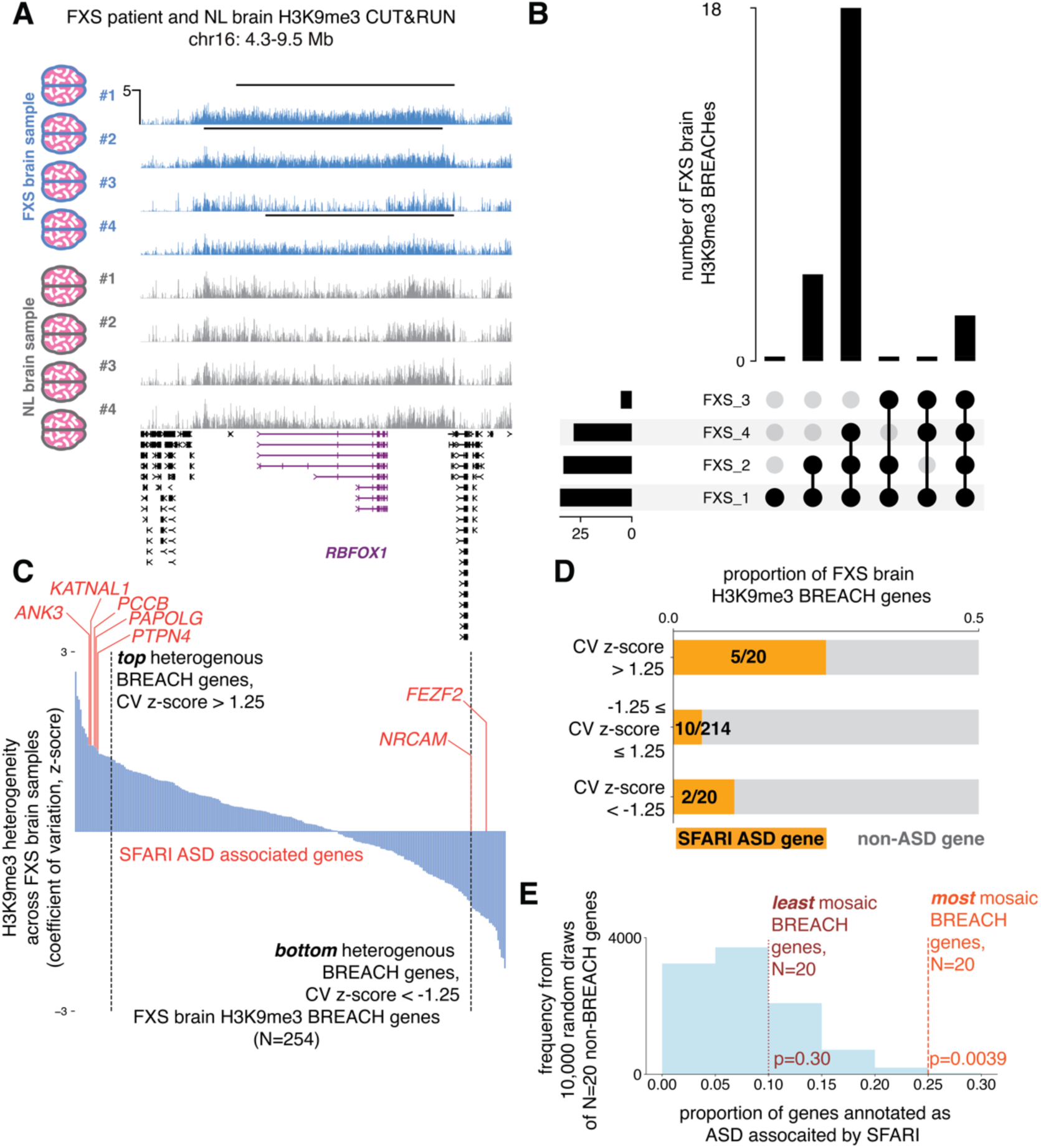
H3K9me3 BREACHes in FXS brain tissue localize with autism-linked genes and exhibit patient-to-patient mosaicism. **(A)** H3K9me3 CUT&RUN from FXS patient brain samples (N=4) and NL donor brain samples (N=4) at a H3K9me3 BREACH on chromosome 16. **(B)** Upset plot with FXS H3K9me3 brain BREACHes (N=40) and the number of FXS brain samples in which they were found. **(C)** FXS H3K9me3 BREACH genes (N=254) ranked by their z-score of coefficients of variation (CV) of H3K9me3 signal across the FXS patient brain samples (N=4). FXS H3K9me3 BREACH genes with highest H3K9me3 heterogeneity across the FXS patient brain samples as defined by a CV z-score > 1.25 that are annotated as autism candidate genes by SFARI are highlighted in red. **(D)** Proportion of FXS brain H3K9me3 BREACH genes with SFARI curated ASD associated genes, stratified by extent of H3K9me3 heterogeneity across FXS patient brain samples. **(E)** Proportion of mosaic and invariant brain BREACH genes (N=20, each) compared to a null distribution of 10,000 draw of random non-BREACH genes (N=20)

### H3K9me3 mosaicism at FXS-specific BREACHes in patient-derived brain tissue and iPSC-derived NPCs

We next set out to classify BREACH genes by the extent to which the H3K9me3 signal is mosaic across FXS patient-derived brain tissue samples. We identified 254 genes colocalized with FXS brain-specific BREACHes (**Table S3**). We formulated a metric to quantify H3K9me3 mosaicism as the coefficient of variation (CV) of H3K9me3 signal in a -2 kb to +10kb window centered on the transcription start site (TSS) for every gene in the N=4 FXS patient brain samples (**Methods**). We ranked BREACH-colocalized genes by their H3K9me3 mosaicism and classified those 1.25 standard deviations or higher above the average as high H3K9me3 mosaic BREACH genes (N=20, **Fig. 1C**) and those with -1.25 standard deviations or less than the average as invariant BREACH genes (N=20, **Fig. 1C**). Our analyses stratify H3K9me3 mosaic and invariant genes colocalized with BREACHes in FXS brain tissue.

To further understand the properties of BREACH genes, we examined their potential association with autism spectrum disorder (ASD). Genes linked to ASD susceptibility as per the SFARI database co-localize with both mosaic and invariant BREACHes (*25*) (**Fig. 1D**). For example, mosaic BREACHes encompass: (1) *ANK3*: a gene on chromosome 10 that encodes an axonal cytoskeleton protein in which missense mutations are associated with intellectual disability, developmental delay, and ASD via a high-confidence SFARI gene score 1 (*26–30*), (2) *KATNAL1*: a gene on chromosome 13 that encodes a microtubule binding protein in which a frameshift mutation is associated with ASD via a strong SFARI gene score 2 (*31–34*), (3) *PCCB*: a gene on chromosome 3 encoding a mitochondrial enzyme implicated in ASD via a high-confidence syndromic SFARI gene score 1S (*35, 36*), (4) *PAPOLG*: an RNA processing gene on chromosome 2 in which de novo missense variants have been associated with ASD via a strong SFARI gene score 2 (*32–34, 37–40*), and (5) *PTPN4*: a gene on chromosome 2 that encodes a protein tyrosine phosphatase implicated in developmental delay, intellectual disability, and ASD (*33, 41–44*). Invariant BREACHes also encompass some genes implicated in ASD via a strong SFARI gene score 2: *FEZF2* (a neural transcription factor gene on chromosome 3 (*26, 45, 46*)) and *NRCAM* (a neuronal adhesion gene on chromosome 7 (*33, 47, 48*)). The proportion of mosaic BREACH genes implicated in ASD is significantly higher than a null distribution of non-BREACH genes genome-wide (**Fig. 1E**). These data suggest that BREACHes are enriched for genes implicated in ASD, intellectual disability, and neurodevelopmental disorders, with mosaic BREACH genes showing stronger confidence SFARI gene scores than invariant BREACH genes and non-BREACH genes.

To ascertain if BREACH mosaicism is unique to brain tissue or also present in FXS patient-derived cell lines, we conducted a similar assessment of H3K9me3 mosaicism in neural progenitor cells (NPCs) derived from induced pluripotent stem cells (iPSC-NPCs). We quantified heterochromatin mosaicism using N=4 FXS patient-derived and N=3 sex-matched neurotypical individual-derived iPSC-NPCs (*24*) (**Fig. 2A**). In addition to our previously published H3K9me3 ChIP-seq libraries (*24*), we generated the same data from a fourth FXS iPSC line (FXS_448) differentiated to iPSC-NPCs using the same established methods (*24*) (**Fig. S1A**). We applied our targeted Nanopore long-read sequencing method, MASTR-seq (*24, 49*), to confirm that the CGG STR in FXS_448 is mutation-length (greater than 200 repeat units) and exhibits classic patterns of DNA hypermethylation over the *FMR1* promoter (*50*) (**Fig. S1B-C**). We generated RNA-seq data from FXS_448 iPSC-NPCs and confirmed the silencing of *FMR1* as previously seen in the other three FXS lines (*24*) (**Fig. S1D**). We compared our *FMR1* promoter DNA methylation data with the *FMR1* RNA-seq data and recapitulated the established anticorrelation between *FMR1* promoter DNA methylation and *FMR1* expression (**Fig. S1E**). Thus, FXS_448 iPSC-NPCs, like the other three previously profiled FXS lines, exhibits the classic molecular features of FXS including a full mutation-length CGG STR tract, DNA methylation at the *FMR1* promoter, and transcriptional silencing of *FMR1*.

**Fig. 2.**
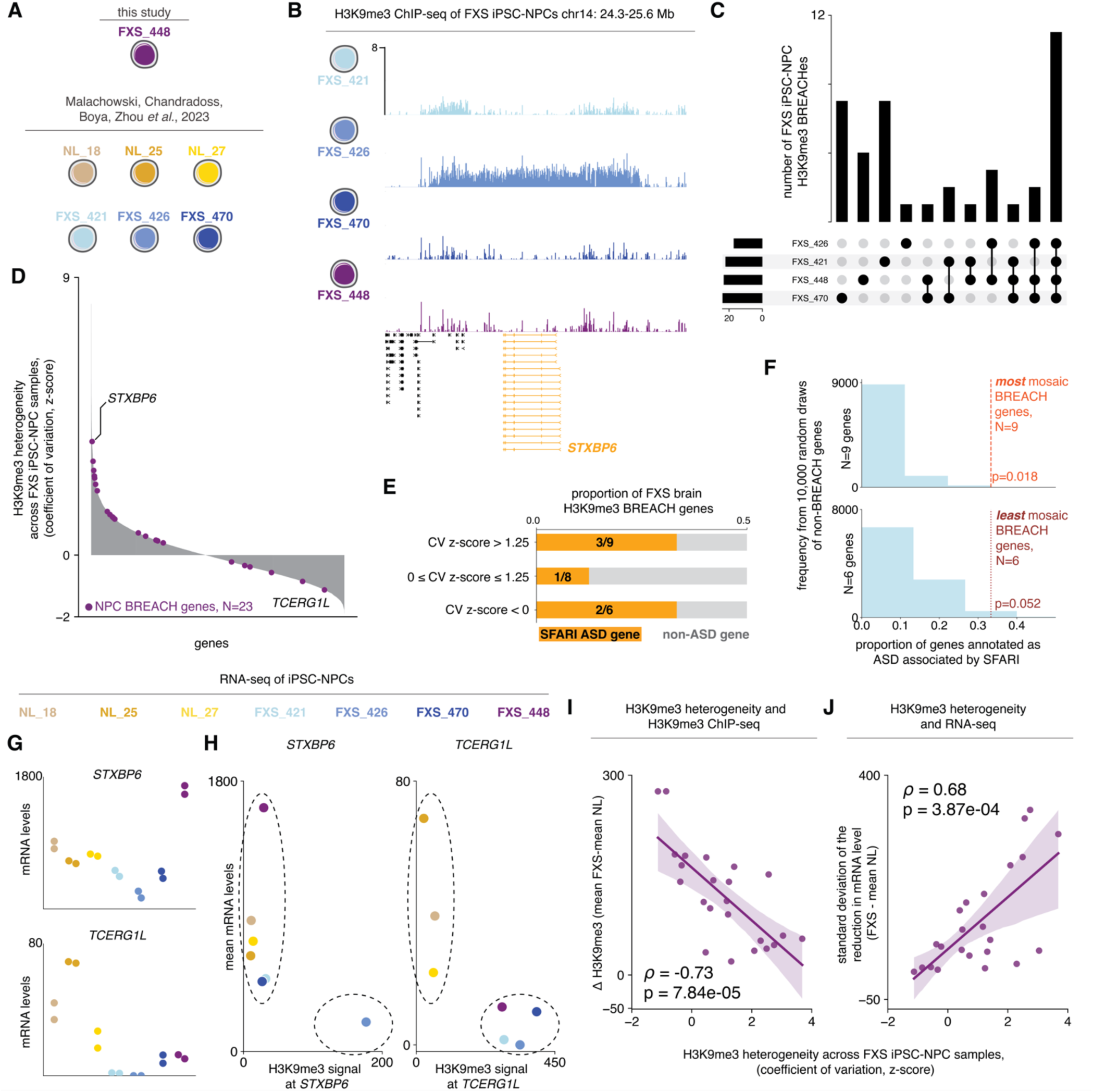
BREACH mosaicism in FXS iPSC-derived NPCs correlates with inter-sample repression of synaptic gene mRNA levels. **(A)** Illustration of a new fourth FXS patient-derived cell line along with the previously characterized three FXS patient-derived cell lines and three NL matched donor-derived cell lines. **(B)** H3K9me3 ChIP-seq from FXS patient iPSC-NPCs (N=4) at a H3K9me3 BREACH on chromosome 14 over *STXBP6*. **(C)** Upset plot of FXS H3K9me3 iPSC-NPC BREACHes (N=40) and the number of FXS iPSC-NPC lines in which they were found. **(D)** Genome-wide ranking of genes by the z-score of coefficients of variation (CV) of H3K9me3 signal across the FXS patient iPSC-NPCs (N=4) with FXS iPSC-NPC BREACH genes in purple dots. **(E)** Proportion of FXS NPC H3K9me3 BREACH genes with SFARI curated ASD associated genes, stratified by extent of H3K9me3 heterogeneity across FXS patient derived iPSC-NPCs. **(F)** Proportion of mosaic (N=9) and invariant (N=6) NPC BREACH genes compared to a null distribution of 10,000 draw of random non-BREACH genes. **(G)** RNA-seq data of *STXBP6* and *TCERG1L* mRNA in the FXS patient iPSC-NPCs (N=4). **(H)** Relationship between H3K9me3 signal at gene promoter of and gene expression across the FXS patient (N=4) and NL matched donor (N=3) iPSC-NPCs, of H3K9me3 heterogeneous NPC BREACH gene, *STXBP6* (left), and H3K9me3 invariant NPC BREACH gene, *TCERG1L* (right). **(I)** Relationship between z-score of H3K9me3 CV across the FXS patient iPSC-NPCs (N=4) and mean difference in H3K9me3 signal between FXS (N=4) and NL iPSC-NPCs (N=3) for each FXS iPSC-NPC BREACH gene (N=23). **(J)** Relationship between z-score of H3K9me3 CV across the FXS patient iPSC-NPCs (N=4) and the standard deviation in the reduction of mRNA levels between FXS (N=4) and the average mRNA levels of NL iPSC-NPCs (N=3) for each FXS iPSC-NPC BREACH gene (N=23).

In our previous publication, we focused on BREACHes with invariant H3K9me3 signal across all FXS iPSC-NPC lines. To understand H3K9me3 mosaicism, we re-classified BREACHes to find N=40 genome-wide present in at least 1 of 4 mutation-length and absent in all 3 normal-length iPSC-NPC lines (**Fig. 2B-C, Table S4, Methods**). We find that nearly 50% of FXS iPSC-NPC BREACHes (N=19/40) were unique to an individual patient-derived iPSC-NPC line, whereas nearly 20% (N=7/40) were found in two, 7.5% (N=3/40) found in three, and nearly 30% (N=11/40) found in all four FXS patient-derived iPSC-NPC lines (**Fig. 2C**). These data confirm that BREACH mosaicism exists in FXS iPSC-NPCs lines as well as brain tissue.

We next classified FXS NPC BREACH genes by the extent of their H3K9me3 mosaicism across FXS iPSC-NPC lines as we did for the brain BREACH genes (**Fig. 2D, Methods**). We used our computed iPSC-NPC H3K9me3 BREACH mosaicism score computed as described above as the coefficient of variation (CV) of H3K9me3 signal in a -2 kb to +10kb window centered on the transcription start site (TSS) for every gene in the N=4 FXS patient iPSC-NPC (**Methods**). We identified 42 genes colocalized with FXS iPSC-NPC BREACHes, 23 of which exhibited detectable mRNA levels using RNA-seq in the iPSC-NPCs (**Table S5**). We stratified BREACH localized genes by their H3K9me3 mosaicism and again observed that the proportion of mosaic BREACH genes implicated in ASD is significantly higher than the null distribution of non-BREACH genes genome-wide (**Fig. 2D-F, Methods**). The highest mosaicism FXS iPSC-NPCs BREACH gene is *STXBP6*, a locus on chromosome 14 involved in neuronal vesical trafficking in which a rare mutation has been described in a patient with developmental epileptic encephalopathy and ASD (*51*) (**Fig. 2D**). Thus, we uncover in both FXS brain tissue and iPSC-NPCs that mosaic and invariant H3K9me3 BREACHes co-localize with genes associated with neurodevelopmental disorders and intellectual disability.

### H3K9me3 mosaicism correlates with inter-sample repression of BREACH genes

H3K9me3 can mark constitutive heterochromatin at pericentromeres, centromeres, and telomeres to repress gene expression and influence genome stability (*52*). H3K9me3 can also form facultative heterochromatin that influences gene expression at lineage-specific genes in a cell type-specific manner (*53*). Therefore, we next set out to ascertain the extent to which BREACH mosaicism is correlated with sample-to-sample variation in gene expression repression. Given the rare nature of FXS patient-derived brain tissue, we elected to dissect the functional role of mosaic BREACH signal on gene expression using iPSC-NPCs. We examined sample-to-sample mRNA level variation using N=4 FXS patient-derived and N=3 sex-matched neurotypical individual-derived iPSC-NPCs. In addition to our recently previously published RNA-seq libraries (*24*), we generated the same data from a fourth FXS iPSC line (FXS_448) differentiated to iPSC-NPCs as previously published (*24*). For the top mosaic BREACH gene, *STXBP6*, we observed wide line-to-line variation in the reduction in mRNA levels that correlates with the variation in H3K9me3 signal (**Fig. 2G-H)**. By contrast, the top invariant BREACH gene, *TCERG1L,* exhibited uniform repression of mRNA levels in FXS iPSC-NPCs (**Fig. 2G-H)**. We note that the hallmark FXS causal gene, *FMR1*, is classified as an invariant BREACH gene with strong and uniform repression of mRNA levels across all FXS iPSC-NPC lines (**Fig. S1D, F**). Our observations provide the foundation for our hypothesis that H3K9me3 mosaicism at BREACHes across different FXS iPSC-NPC lines is linked to line-specific heterogeneity in the reduction of mRNA levels per line.

Beyond locus-specific examples, we sought to assess the relationship between the extent of sample-to-sample variation in gene expression repression and mosaicism in H3K9me3 signal at BREACH co-localized genes. To verify that our H3K9me3 mosaic metric strongly correlates with bulk levels of H3K9me3 signal, we confirmed the expected strong anti-correlation between the extent of mosaicism of H3K9me3 at each gene and the average change in H3K9me3 signal across FXS compared to NL iPSCs (**Fig. 2I**). We also compared each BREACH gene’s H3K9me3 mosaicism score to the standard deviation in the reduction in mRNA level across the N=4 FXS iPSC-NPC lines compared to the mean of the N=3 normal-length iPSC-NPC lines. We identified a strong correlation between the H3K9me3 mosaic score in FXS iPSC-NPCs and the sample-to-sample variation in the reduction in mRNA levels between each of the N=4 FXS iPSC-NPC lines and the average of the N=3 normal-length iPSC-NPC lines (**Fig. 2J**). Given the strong observed relationships are calculated in population-based RNA-seq and ChIP-seq, we hypothesize that the bulk observations reflect high heterogeneity in mRNA levels across lines and across single-cells. Together, our data demonstrate that the variation in gene expression repression in FXS iPSC-NPCs is correlated with the mosaicism of H3K9me3 signal at BREACHs across individual patient donors in FXS.

### Link between H3K9me3 mosaicism and heterogeneity in 3D genome miswiring, CTCF occupancy, and mRNA levels at the FMR1 BREACH in FXS patient-derived iPSC-NPCs

We previously reported that H3K9me3 at BREACHes localizes with the dissolution of higher-order genome folding patterns of topologically associated domains (TADs), subTADs, and long-range loops as well as loss of occupancy of the architectural protein CCCTC-binding factor (CTCF) (*24*). Given that our new FXS_448 iPSC-NPC line shows variable expression of BREACH genes like the other FXS iPSC-NPC lines (**Figure 2**), we created Hi-C and CTCF ChIP-seq data for FXS_448 iPSC-NPCs (**Figure 3**). We confirm that there is no detectable H3K9me3 domain and intact 3D genome folding and CTCF occupancy patterns in all sex-matched control NL_18, NL_25, and NL_27 iPSC-NPCs as previously reported (**Fig. S2A**). As previously reported using FXS_421, FXS_426, and FXS_470 (*24*), we find an H3K9me3 BREACH on chromosome X spanning *FMR1* which can spread up to 8 Mb upstream in an FXS iPSC-NPC line specific manner to variably cover two neuronal adhesion genes, *SLITRK4* and *SLITRK2* (**Fig. 3A, Fig. S2A**). Importantly, we see that the H3K9me3 in FXS_448 iPSC-NPCs is substantially lower in magnitude and spread compared to the other three FXS iPSC-NPC lines (**Fig. 3A).** Specifically, H3K9me3 signal in FXS_448 iPSC-NPCs is lower in magnitude compared to the other 3 FXS iPSC-NPC lines over *SLITRK2* (**Fig. 3A and 3E**), and it fails to spread to encompass *SLITRK4* just as with FXS_426 and FXS_470 (**Fig. 3A-B**). Severe disruptions in 3D genome folding and loss of CTCF occupancy present in FXS_421, FXS_426, and FXS_470 do not occur in FXS_448 in direct correlation with the low H3K9me3 signal. Together, these results indicate that the extent of the mosaicism in the spread of the H3K9me3 signal correlates with the genomic range in which TADs, subTADs, loops, and CTCF binding sites are abolished in FXS iPSC-NPCs.

**Fig. 3.**
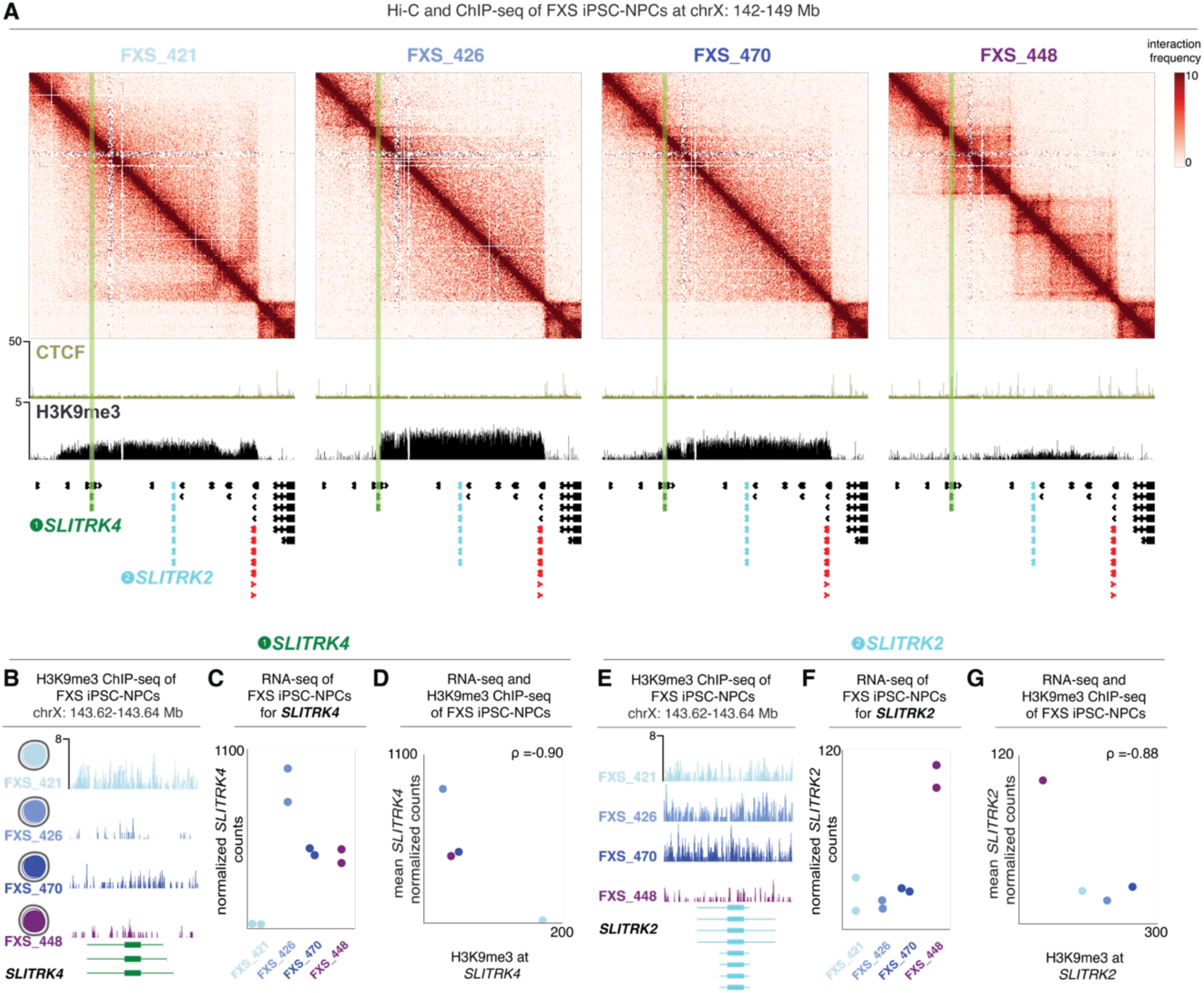
Variation in heterochromatin deposition correlates with line-to-line differences in higher-order genome misfolding and reduction in mRNA levels across FXS patient-derived iPSC-NPCs. **(A)** Hi-C heatmaps and CTCF and H3K9me3 ChIP-seq tracks of FXS patient iPSC-NPCs at a 7 Mb region on the X chromosome including *SLITRK4* (green), *SLITRK2* (blue), and *FMR1* (red). Green highlight over *SLITRK4*. **(B)** H3K9me3 ChIP-seq over *SLITRK4* in the FXS patient iPSC-NPCs (N=4). **(C)** RNA-seq for *SLITRK4* mRNA in the FXS patient iPSC-NPCs (N=4). **(D)** Relationship between H3K9me3 signal at *SLITRK4* and *SLITRK4* expression across in FXS patient iPSC-NPCs (N=4). **(E)** H3K9me3 ChIP-seq over *SLITRK2* in the FXS patient iPSC-NPCs (N=4). **(F)** RNA-seq for *SLITRK2* mRNA in the FXS patient iPSC-NPCs (N=4). **(G)** Relationship between H3K9me3 signal at *SLITRK2* and *SLITRK2* expression across the FXS patient iPSC-NPCs (N=4).

To assess the link between H3K9me3 mosaicism and gene expression at the X chromosome BREACH, we examined RNA-seq and H3K9me3 ChIP-seq signal at *FMR1*, *SLITRK4*, and *SLITRK2*. The X chromosome BREACH spreads over *SLITRK4* only in FXS_421 iPSC-NPCs (**Fig. 3B**) and only the FXS line exhibits repressed mRNA levels of *SLITRK4* (**Fig. 3C**). Moreover, the X chromosome BREACH shows strong signal over *SLITRK2* in FXS_421, FXS_426, and FXS_470 but is negligible in FXS_448 iPSC-NPCs (**Fig. 3E**) and only FXS_448 iPSC-NPCs exhibit high expression of *SLITRK2* mRNA levels (**Fig. 3F**). We find that mRNA levels strongly anti-correlate with H3K9me3 signal among the four FXS patient iPSC-NPCs (**Fig. 3D and G**). Our data suggest a strong correlation between H3K9me3 and the dissolution of CTCF binding, 3D genome folding patterns, and reduced gene expression in FXS patient-derived iPSC-NPCs.

### H3K9me3 mosaicism across isogenic subclones from a single FXS patient-derived iPSC-NPC line correlates with the severity of gene expression repression

Since the four FXS patient iPSC-NPCs were derived from different individuals, *trans* modifiers in the genetic background could confound correlations between H3K9me3 and mRNA levels. To rule out differences in genetic background, we re-analyzed H3K9me3 CUT&RUN data (*24*) published in N=7 subclones from the FXS_421 iPSC line derived from a singular FXS patient (**Fig. 4A**). We conducted qRT-PCR for *SLITRK2* and *SLITRK4* in the FXS_421 iPSC subclones and observed substantial heterogeneous mRNA expression of *SLITRK4* and *SLITKR2* (**Fig. 4B-C**). With H3K9me3 CUT&RUN, we observed that the H3K9me3 signal at the BREACH variably extends over *SLITRK2* and *SLITRK4* (**Fig. 4D-E**). We observed a strong anti-correlation between H3K9me3 signal at *SLITRK4* and *SLITRK2* mRNA levels in the isogenic FXS iPSC subclones (**Fig. 4F**). These data demonstrate the correlation between H3K9me3 signal and the extent of the reduction in mRNA levels across single-cell clones derived from the same FXS genetic background.

**Fig. 4.**
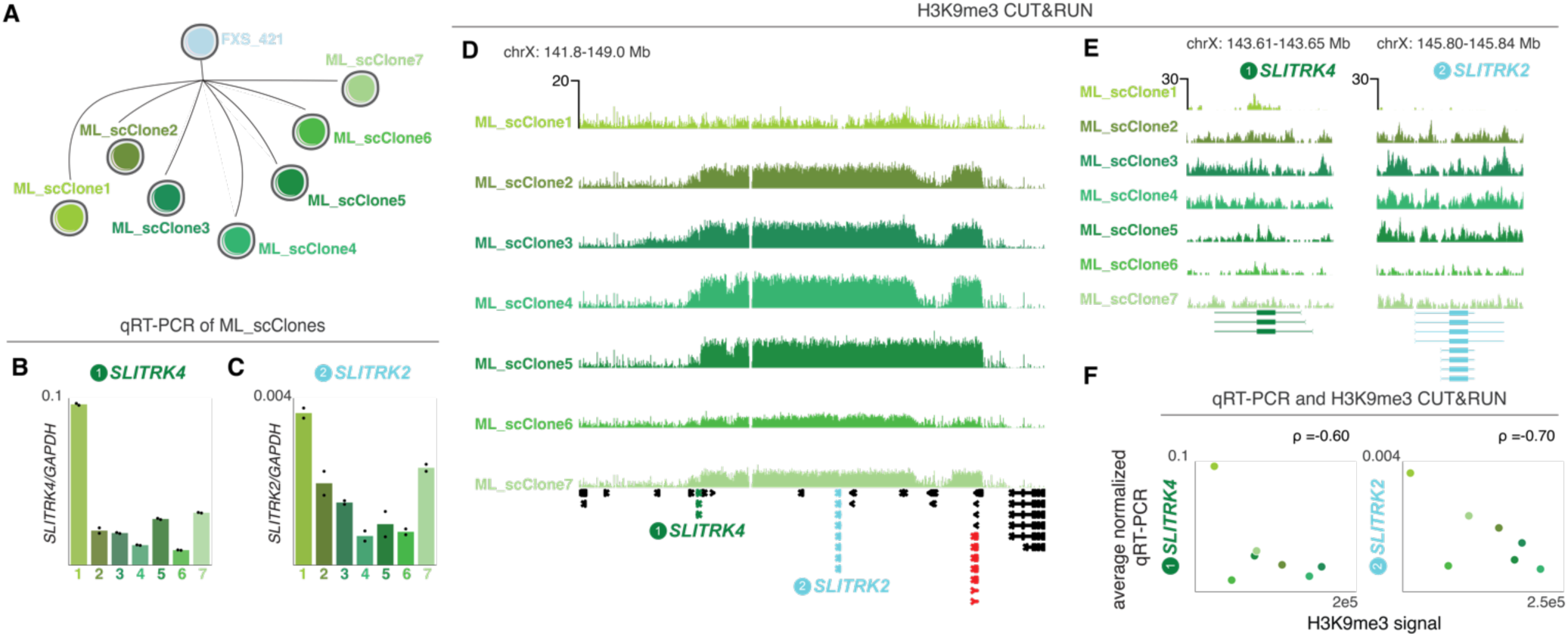
Heterochromatin deposition anti-correlates with synaptic gene expression across subclones from a single FXS patient-derived parent iPSC line. **(A)** Schematic of seven previously characterized subclones of parental FXS_421 (*24*) **(B-C)** Normalized qRT-PCR (N=2) of (B) *SLITRK4* and (C) *SLITRK2* mRNA levels normalized by *GADPH* mRNA level. **(D-E)** H3K9me3 CUT&RUN over (D) 7 Mb on the X chromosome (141.8-149.0 Mb), (E) *SLITRK4* and SLITRK2 in the seven ML_scClones (24). **(F)** Relationship between H3K9me3 signal at gene promoter of and gene expression across the subclones of parental FXS_421 (N=7) of *SLITRK4* (left) and *SLITRK2* (right).

### mRNA levels of H3K9me3-mosaic genes increase the predictive power of FMRP levels for clinical metrics of adaptive functioning and cognitive ability in FXS patients

FMRP levels in blood samples are a molecular correlate for FXS severity (*20*). We assessed the predictive value of mosaic H3K9me3 and invariant H3K9me3 BREACH genes on clinical measures of adaptive functioning and cognitive ability in FXS patients. We used a published dataset of N=16 sex-matched (male) FXS patients that includes molecular profiling of gene expression and FMRP levels in blood samples and a clinical measurement of adaptive functioning called the Vineland-3 Adaptive Behavior Composite (ABC) score in the same patients (*54*). The ABC score is used as part of the NIH Toolbox Cognition Battery in the longitudinal assessment of FXS patients with the aim of using of these tools as outcome measurements for treatment trials (*55*). Although the sample size is small, this is the one available dataset we are aware of that allows for an assessment of the ability of BREACH genes to add predictive value to FMRP levels for the identification of novel molecular correlates of FXS severity.

We built an additive model of ABC score as the dependent variable and FMRP as the independent variable after first confirming the expected significant correlation between ABC score and FMRP levels as previously reported (Pearson’s ρ = 0.47, **Fig. 5A**) (*20, 54*). FMRP only explains 22% of the variance of the ABC score and does not significantly contribute to the model (q-value = 0.09), highlighting the need for additional biomarkers that can be assessed in the clinic (**Fig. 5A**). As a discovery set, we first start with the N=42 genes co-localized with BREACHes in FXS patient-derived iPSC-NPCs (**Figure 2**). We filtered for genes encoding mRNA with detectable transcripts in FXS patient blood. We added each gene individually to an additive model and computed a p-value and a q-value to assess improvements in ABC predictive power (**Fig. 5B, Methods, Table S6**). Of the mRNA-encoding detectable genes localized with FXS iPSC-NPC BREACHes, we classified 7 genes as H3K9me3 mosaic (*AJAP1, IL5RA, KHDRBS2, SLITRK4, STXBP6, TRNT1, VIPR2*), and 4 genes as H3K9me3 invariant (*FKBP1C, GLT1D, GZMH, SLC15A4*) (**Fig. 5C**). We performed the same analysis for all detectable non-BREACH genes genome-wide (N=13992) and observed that the mosaic genes exhibited more predictive power in an additive model with FMRP for ABC compared to non-BREACH control genes (**Fig. 5D**). The four invariant BREACH genes did not add predictive power to the ABC prediction (**Fig. 5E**). The two strongest contributors to the ABC predictive model were *SLITRK4* and *KHDRBS2* (**Fig. 5C**) with significant q-values compared to a model of FMRP alone. Both *SLITRK4* and *KHDRBS2* are mosaic genes. We repeated these analyses with brain BREACH genes and found that only one brain BREACH gene, the mosaic BREACH gene, *ANGEL2,* improved the model with ABC over FMRP alone (**Fig. S3A-B**). Thus, the mRNA levels of select mosaic BREACH genes in blood measurements adds explanatory power compared to FMRP alone in predicting ABC scores in FXS patients.

**Fig. 5.**
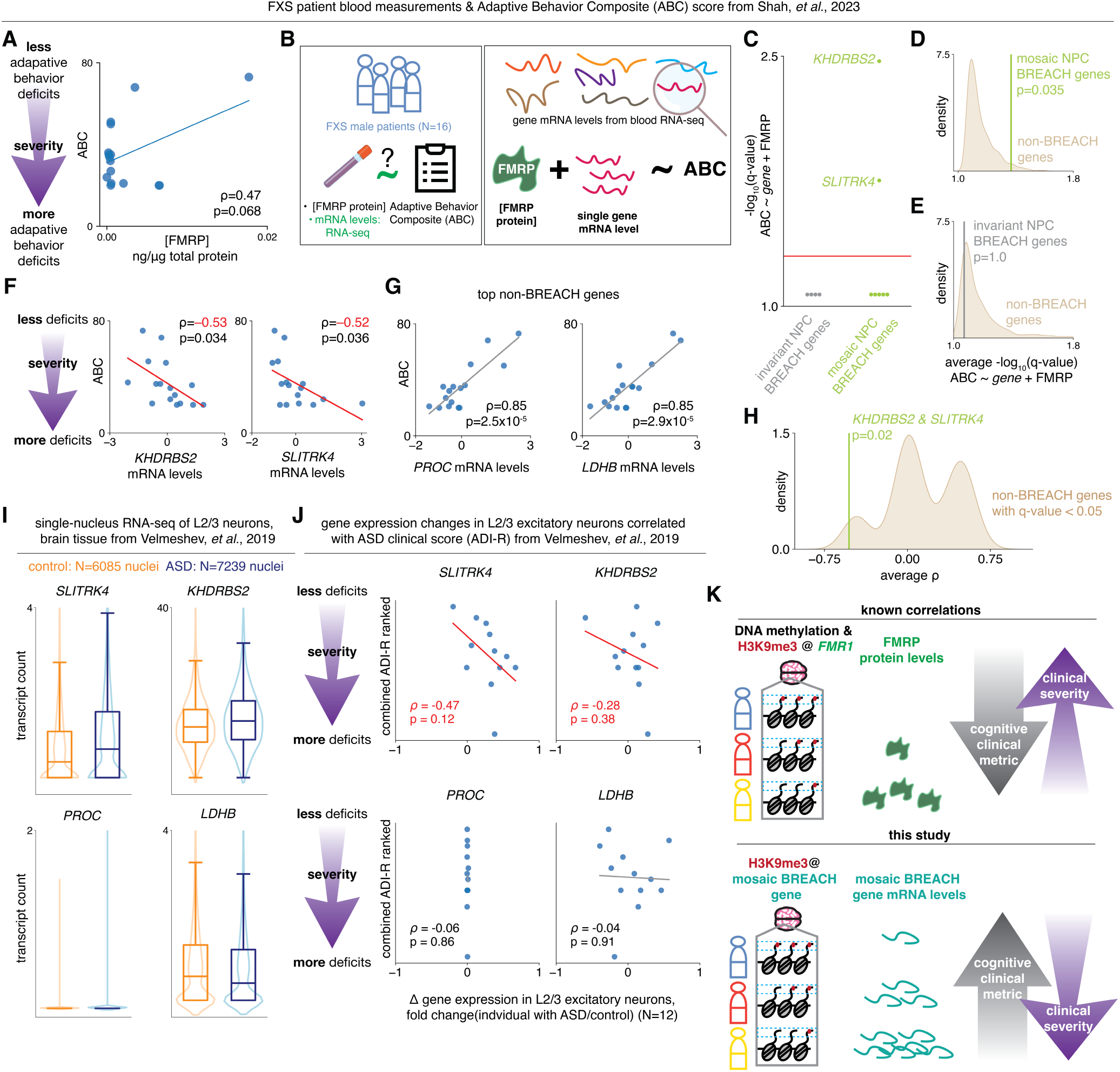
Mosaic FXS BREACH genes mRNA in FXS patient blood is associated with greater clinical severity in FXS patients unlike classic FXS prognostic, FMRP. **(A)** Relationship between FXS patient blood FMRP protein level and measure of Adaptive Behavior Composite, ABC, across FXS patients (N=16) (*54*). **(B)** Schematic of search for genes whose mRNA level in FXS patient blood alongside FMRP protein levels in FXS patient blood predict Adaptive Behavior Composite (ABC) score in an additive model. **(C)** Q-values for the correlation of the pair of an individual gene’s FXS patient blood mRNA level and FMRP protein level with ABC score for mosaic and invariant NPC BREACH genes. **(D-E)** Average -log10(q-value) of **(D)** mosaic (N=7) and **(E)** invariant (N=4) NPC BREACH genes compared to a null distribution of 1000 draw of random non-BREACH genes (matched N per comparison). **(F)** Relationship between FXS patient blood mRNA levels of the most predictive mosaic NPC BREACH genes, *KHDRBS2* or *SLITRK4,* and measure of Adaptive Behavior Composite, ABC, across FXS patients (N=16). **(G)** Relationship between FXS patient blood mRNA levels of the most predictive non-NPC-BREACH genes, *PROC* or *LDHB,* and measure of Adaptive Behavior Composite, ABC, across FXS patients (N=16). **(H)** Average rho of mosaic BREACH genes with a q-value less than 0.05 (*KHDRBS2* and *SLITRK4*) compared to a null distribution of 1000 draw of random non-BREACH genes with a q-value < 0.05. **(I)** Single-nuclei RNA-seq expression from L2/3 excitatory neurons from ASD patient and control brains (*60*) for the most predictive mosaic NPC BREACH genes, *KHDRBS2* or *SLITRK4*, and most predictive non-NPC-BREACH genes, *PROC* or *LDHB*. **(J)** Relationship between ASD patient mRNA levels of the most predictive mosaic NPC BREACH genes, *KHDRBS2* or *SLITRK4*, and most predictive non-NPC-BREACH genes, *PROC* or *LDHB*, and combined Autism Diagnostic Interview–Revised (ADI-R) score ranks (N=12) (*60*). **(K)** DNA methylation and H3K9me3 are anticorrelated with FMRP levels in FXS patients. At mosaic BREACH genes, variation of H3K9me3 across FXS patients explains variation in gene expression. Levels of the classic clinical correlate in FXS, FMRP, are associated with higher clinical metrics and less severe presentations. However, the mRNA levels of mosaic BREACH genes are associated with lower clinical metrics and greater clinical severity in patient blood.

To understand the relationship between mosaic BREACH gene mRNA levels in FXS patient blood and ABC, we correlated each gene’s mRNA level with ABC score. In the case of FMRP, we observed the long-time known correlation between protein levels and ABC score, thus confirming that low FMRP occurs with low ABC and therefore decreased cognitive functioning in disease (**Fig. 5C**). We unexpectedly found that mRNA levels of *SLITRK4*, *KHDRBS2*, and *ANGEL2* were anti-correlated with FXS severity (**Fig. 5F, Fig. S3B**). By contrast to FMRP, all three mosaic BREACH genes exhibited decreased ABC score with high mRNA levels and, hence, higher gene expression occurs with low ABC and decreased cognitive performance in FXS (**Fig. 5F**). The top non-BREACH genes that significantly contribute to the additive model have positive slopes indicating that higher expression correlates with higher adaptive functioning and hence a higher ABC metric (**Fig. 5G**). Using a resampling test, we demonstrate that the mosaic BREACH genes are significantly more likely to contribute to our ABC score predictive model through with a negative slope compared to non-BREACH genes that tend to show positive slopes (**Fig. 5H**). Our analyses reveal that mosaic BREACHes across FXS iPSC-NPCs can help nominate unexpected genes, in this case *SLITRK4* and *KHDRBS2*, which improve the predictive power of FMRP for adaptive functioning via an anti-correlation with the ABC score.

We then assessed possible secondary correlations between the predictive mosaic BREACH genes and the established molecular correlates of FXS: (1) size mosaicism (alleles with *FMR1* CGG STRs of repeat length less than 200 (*56–58*)) and (2) methylation mosaicism (alleles with unmethylated *FMR1* promoters (*15–19*)). We found that *SLITRK4*, *KHDRBS2*, and *ANGEL2* mRNA levels did not differ significantly between FXS patient samples with and without size mosaicism or between FXS patients with and without alleles with unmethylated *FMR1* promoters (**Fig. S4**). Thus, we did not find evidence that mosaicism in previously reported molecular correlates of FXS could explain additional predictive power for the ABC clinical metric from the mosaic BREACH genes.

To validate the anti-correlation between mosaic BREACH genes and clinical adaptive functioning metrics we re-analyzed data from an independent study. Individuals with FXS are commonly co-diagnosed with ASD, and FXS is the monogenic cause of ASD (*59*). We re-analyzed single-nucleus RNA-seq of cortical tissue from individuals with and without a diagnosis of ASD in which gene expression changes from the L2/3 cortico-cortical projection neurons were reported as most correlated with clinical measurements of ASD symptom severity (*60*). Consistent with our FXS patient blood data (**Fig. 5F-H**), we find that *SLITRK4* and *KHDRBS2* showed increased expression in L2/3 excitatory neuronal nuclei from ASD donors compared to the control donors (**Fig. 5I**). In contrast, the top non-BREACH genes were minimally expressed and show no difference in expression (**Fig. 5I**). Finally, we correlated the expression of the predictive mosaic BREACH genes and the predictive non-BREACH genes with a composite rank based on the Autism Diagnostic Interview–Revised (ADI-R) clinical metric in which a higher rank corresponds to less deficits. We find that all three predictive mosaic BREACH genes show a negative correlation with the combined ADI-R rank just as with the ABC score (**Fig. 5J, S3C**). Altogether, our data highlight a working model in which mosaic BREACH genes improve the predictive power of FMRP for physiologically relevant clinical metrics of adaptive functioning and cognition in FXS patients (**Fig. 5K**).

## Discussion

We recently reported the Megabase-scale domains of the heterochromatic histone modification H3K9me3 in FXS patient-derived iPSCs and iPSC-NPCs (*24*). However, the variation of BREACHes in FXS patient-derived brain tissue and the potential clinical relevance of these heterochromatin domains is unknown. Applying histone profiling, 3D genome folding, and genome-wide gene expression assays to rare FXS patient-derived brain samples, FXS patient-derived iPSC-NPC lines, and subclones from a single FXS iPSC line, we demonstrate a strong correlation between mosaic inter-sample H3K9me3 signal and heterogeneous mRNA levels. Altogether, our data leverages the variability of H3K9me3 at BREACHes across FXS patient samples to demonstrate the association between H3K9me3 and repression of BREACH co-localized genes, which are associated with neurodevelopmental disorders, intellectual disability, neurogenesis, and neuronal function. We demonstrate improved prediction of clinical metrics of adaptive functioning and cognition in FXS patients with an additive linear model accounting for blood measurements of FMRP and mRNA levels of BREACH-mosaic genes compared to the current standard of FMRP levels alone. Together, our data suggest that H3K9me3 mosaicism at BREACHes can identify and prioritize genes with patient-to-patient mosaic repression that have predictive value for cognitive metrics in FXS patients (**Fig. 5K**).

FXS patients presents with clinical heterogeneity, underscoring the need to better understand the sources of variation across patients to better understand the pathophysiology and enable therapeutic insights for precision-medicine based approaches (*61*). FXS patients with unmethylated *FMR1* promoter despite a full mutation-length CGG STR had higher levels of FMRP as measured in their blood, which predicted less severe clinical presentations (*15, 20–22*). These foundational studies and observations demonstrate that epigenetic mosaicism is associated with the severity of FXS clinical presentations and highlight the power of blood-based profiling to aid in elucidating the link between epigenetic mosaicism and disease severity. Here, our work extends the paradigm by analyzing a dataset of FXS male patients with blood RNA-seq, FMRP levels, and ABC score measurements. We find the addition of mosaic FXS BREACH genes improves the correlation beyond the classic molecular correlate – FMRP levels – of FXS clinical severity (*20*). While FMRP levels are associated with better performance on clinical metrics (*20, 54*), we find that the blood levels of the mosaic FXS BREACH genes (*SLITRK4* and *KHDRBS2* and *ANGEL2*) are anti-correlated with cognitive clinical metrics. By contrast, non-FXS BREACH genes tend to behave like FMRP levels and correlate with cognitive clinical metrics. To validate this surprising finding, we analyzed single-cell genomics data paired with clinical metrics relevant to autism and confirm the anti-correlation between BREACH-mosaic genes, *SLITRK4*, *KHDRBS2*, and *ANGEL2,* and an autism cognitive score. Thus, we uncover that the expression of mosaic BREACH genes is a source of molecular variation across patients that are anti-correlates of cognitive clinical scores relevant to FXS and autism.

Higher levels of FMRP are associated with less severe FXS symptoms (*15, 20–22*) (**Fig. 5K**, *top*). Loss of FMRP in FXS leads to impaired mRNA splicing, dysregulation of translation, and dysfunction of other proteins that FMRP binds (*62*). Molecular deficits in proper FMRP function contributes to defects in neuronal development, axonal targeting, and synaptic connectivity and function (*63, 64*). A leading hypothesis of FXS pathophysiology suggests an increase in neuronal excitability can manifest as FXS symptoms, such as social anxiety and hypersensitivity to stimulus (*63, 65*). Moreover, neuronal hyperexcitability—characterized by increased firing rates and burst frequency *in vitro*—has been rescued in FXS iPSC-derived neurons through *FMR1* reactivation (*66–68*). Thus, escape from repression and higher levels of FMRP is associated with rescue of neuronal function and better clinical metrics in FXS patients.

Here, in the present study, the FMRP results in our data is consistent with published literature. Using FXS patient derived data of cognitive metrics and molecular protein and mRNA levels from blood samples, we generated an additive model of blood FMRP levels combined with gene expression and clinical metrics. We demonstrate that FMRP alone significantly contributes to prediction of a clinical ABC score. The expected positive slope confirms the known relationship of higher FMRP levels resulting in higher cognitive ability (*15, 20–22*). Thus, FMRP alone in our data shows a moderate but positive correlation to cognition in FXS patients.

By contrast to FMRP, higher levels of mosaic BREACH mRNAs anti-correlate with the ABC score and thus higher disease severity (**Fig. 5K**, *bottom*). A potential mechanism through which the higher levels of mosaic BREACH genes could mediate more severe FXS presentation is disrupting neuronal excitability in FXS. While the role for *ANGEL2* in neuronal function is unknown, *SLITRK4* and *KHDRBS2* play pivotal roles in synaptogenesis (*69–72*). First, *SLITRK4* is a neuronal adhesion gene that has been previously associated with neurodevelopmental disorders (*73*). Increasing *Slitrk4* levels in cultured mouse hippocampal neurons resulted in an increase in excitatory synapse density (*70*). Second, *KHDRBS2* encodes for an RNA-binding protein that regulates the splicing of neurexins, neuronal adhesion molecules with significant associations to neurodevelopmental disease (*72, 74–76*). Overexpression of *Khdrbs2* mis-splices neurexin isoforms, altering its ligand affinities and increasing network activity in mouse models (*72, 77, 78*). We hypothesize based on our early data here that higher expression of the mosaic BREACH genes *SLITRK4* and *KHDRBS2* in FXS patients could contribute to the disruption of synaptic plasticity, synapse development, or neurophysiology. Our working model is that individuals with lower H3K9me3 would have higher expression of mosaic BREACH genes *SLITRK4, KHDRBS2,* and *ANGEL2*, which could contribute to severe FXS presentations by exacerbating synaptic or neurophysiological defects. Future work should aim to understand the potential functional role of mRNA levels of BREACH mosaic genes in normal-length, neurotypical individuals as well as a range of expression in FXS patients and their connection to neural function.

Regulation of mosaic FXS BREACH synaptic gene expression by H3K9me3 reflects the contribution of chromatin regulation in FXS (*24, 63, 79*). Additionally variants in enzymes responsible for the deposition and removal of H3K9me3 have been identified in syndromic forms of autism spectrum disorder (*37, 46, 80–83*) and in patients with neurodevelopmental delay (*84*), demonstrating the influence of H3K9me3 in neurodevelopment and neuronal function. Our analyses here of FXS patient samples support heterochromatin as a regulator of gene expression and contributor to FXS clinical presentations, raising the potential of targeting chromatin-modifying enzymes as therapeutic strategies to ameliorate synaptic and neurophysiological pathologies in FXS.

## Materials and Methods

Some of the methods described below have been detailed in our own previous manuscripts (*24, 85–101*). We describe the methodological steps below even in cases where we have published them before to ensure rigor and reproducibility.

### Donor brain tissue

We used NIH NeuroBioBank post-mortem caudate nucleus brain tissue from male donors with a clinical diagnosis of fragile X syndrome (FXS) and male donors without a clinical diagnosis of FXS (donor age, sex, ethnicity and race in **Table S1**). We kept brain tissue at -80°C.

### Fluorescence-activated nuclei sorting on donor brain tissue

We processed the post-mortem caudate nucleus brain tissue as previously described to enable harmonized comparison across donor samples (*24*). We sectioned the tissue into ∼100 mg fragments using sterile forceps and single use razor blade, working on dry ice. We then proceeded to continue to process approximately 100 mg of caudate nucleus at a time from each donor, working on ice. We added the tissue into 10 mL of ice-cold Homogenization Buffer (0.32 M sucrose (Sigma-Aldrich, S0389-500G), 5 mM CaCl_2_ (Thermo Fisher, J63122-AD), 10 mM Tris-HCl pH 8.0 (Invitrogen, 15568025), 3 mM MgAc_2_ (Sigma-Aldrich, 63052-100ML), 0.1% Triton X-100 (Sigma-Aldrich, T8787-100ML), 0.1 mM EDTA (Invitrogen, 15575020), 1x Protease Inhibitor Cocktail (Roche, 11873580001)). We used the 15 mL Dounce Tissue Grinder (Wheaton, 357544) to dounce the tissue with 20 strokes of the loose pestle and 7 strokes of the tight pestle on ice. We transferred the dounced solution onto 14 mL of ice cold sucrose cushion (1.8 M sucrose (Sigma-Aldrich, S0389-500G), 10 mM Tris-HCl pH 8.0 (Invitrogen, 15568025), 3 mM MgAc_2_ (Sigma-Aldrich, 63052-100ML), 1x Protease Inhibitor Cocktail (Roche, 11873580001)) and added 12 mL ice cold Homogenization Buffer on top of the dounced solution and centrifuged at 25,700 RPM (∼81k xg, SW Ti 32 rotor) for 2 hours in a swinging bucket rotor in a centrifuge at 4°C.

We removed the supernatant and added ice cold FANS Buffer (1x PBS (Corning, 21-040-CV), 1% bovine serum albumin (Sigma-Aldrich, A7906-50G), 1x Protease Inhibitor Cocktail (Roche, 11873580001)). We incubated the pellet on ice for 20 mins, then resuspended and performed a nuclei count. After centrifugation of the solution at 600xg for 6 mins at 4°C, we added FANS buffer to a concentration of 3 million nuclei/mL and rotated the solution for 15 mins at 4°C for blocking. While rotating at 4°C, we added Alexa Fluor 488 conjugated anti-NeuN (1:1000, Sigma-Aldrich, MAB377X) for 90 mins and DAPI (1:2000, Sigma-Aldrich, MBD0015-1ML) for the final 5 mins. After centrifugation for 6 mins at 4°C at 600xg, we resuspended nuclei in FANS buffer for a final concentration of 6 million nuclei/mL, filtered into a 5 mL FACS sorting tube (Corning, 352235), and sorted using the MoFlo Astrios (Beckman Coulter).

### CUT&RUN on donor brain tissue

We performed CUT&RUN (*24, 102*) with the recovered nuclei into CUT&RUN Wash Buffer (20 mM HEPES-KOH pH 7.5 (Boston BioProducts, BBH-75-K), 150 mM NaCl (Invitrogen, AM9760G), 0.5 mM Spermidine (Sigma-Aldrich, S2501), 1x Protease Inhibitor Cocktail (Roche, 11873580001)) and immediately bound nuclei to Concanavalin A beads (BioMag, 86057). We resuspended the bead-bound nuclei in 100 μL antibody buffer (CUT&RUN Wash Buffer with 0.1% Triton X-100 (Sigma-Aldrich, 93443) and 2 mM EDTA (Invitrogen, 15575020)) and added 1 μg of antibody (IgG (Sigma-Aldrich, I8140) or H3K9me3 (Abcam, ab8898)). We incubated the solution rotating overnight at 4°C to allow for antibody binding.

The next day, after 3 washes with Triton Wash Buffer (Wash Buffer with 0.1% Triton X-100), we resuspended the bead-bound nuclei in 50 μL Triton Wash Buffer and added 2.5 μL of CUTANA pAG-MNase (EpiCypher, 15-1016). After 10 minutes at room temperature, we washed twice with Triton Wash Buffer, resuspended the nuclei with 100 μL Triton Wash Buffer. After 5 minutes on ice, we added 2 μL of 100 μM CaCl_2_ (Thermo Fisher, J63122-AD). After 2 hours of incubation with rotation at 4°C for MNase digestion, we added 100 μL of 2X Stop Buffer (340 mM NaCl, 20 mM EDTA, 4 mM EGTA (BioWorld, 40520008-1), 0.04% Triton X-100 (Sigma-Aldrich, 93443), 50 μg/mL RNase A (Thermo Scientific, EN0531), 50 μg/mL Glycogen (Thermo Scientific, R0561)) and incubated the nuclei at 37°C for 30 minutes to allow for chromatin release. After magnetic immobilization of the bead bound nuclei, we collected the supernatant and performed phenol:chloroform extraction and ethanol precipitation of the DNA. We used the NEBNext Ultra II Library Prep Kit (New England Biolabs, E7645S) per the manufacturer’s instructions. We sequenced CUT&RUN libraries with 37 bp paired end reads on an Illumina NextSeq 500.

### Induced pluripotent stem cell (iPSC) culture

We used an iPSC line derived from a male FXS patient, GM09497, designated FXS_448 in this study). We received the iPSC line from Fulcrum Therapeutics, where the line was expanded and characterized. We additionally used seven previously characterized subclones of parental FXS_421 iPSCs, ML_scClones1-7 and cultured them as previously described (*24, 103*). Briefly, we cultured iPSCs on plates coated with Matrigel hESC-Qualified Matrix (Corning, 354277) at 37°C and 5% CO_2_. We used mTeSR™ Plus media (STEMCELL Technology, 100-0276) supplemented with 1% penicillin/streptomycin (Gibco, 15140122). Every 2-4 days, we passaged iPSC that reached 70-80% confluence.

### Neural progenitor cell (NPC) induction from iPSCs

We differentiated iPSCs to NPCs as previously described (*24, 104*). We dissociated iPSCs from culture plates with Accutase (Gibco, A1110501). After washing out the Accutase, we replated them at a density of 1e5 cells per cm^2^ in NPC differentiation media on plates coated with Matrigel hESC-Qualified Matrix (Corning, 354277). NPC differentiation media included 5 ug/mL insulin (Sigma-Aldrich, I1882), 64 mg/mL L-ascorbic acid (Sigma-Aldrich, A8960), 14 ng/mL sodium selenite (Sigma-Aldrich, S5261), 10.7 ug/mL Holo-transferrin (Sigma-Aldrich, T0665), 543 mg/mL sodium bicarbonate (Sigma-Aldrich, S5761), 10 mM SB431542 (STEMCELL Technology, 72234), and 100 ng/mL Noggin (R&D Systems, 6057-NG) in DMEM/F-12 (Gibco, 11320033). After eight days of culture with daily media changes, we assessed them for rosette structure and harvested the iPSC-NPCs.

### MASTR-seq: targeted Nanopore long-read sequencing of the *FMR1* locus

We performed MASTR-seq as previously described (*24, 49*). We refer to the MASTR-seq manuscript for a detailed step-by-step protocol (*105*). First, we prepared high molecular weight DNA preparation from ∼ 5 million iPSCs. We lysed the cells in a 10 ML solution of 10 mM Tris-Cl (pH 8) Tris-Lysis-Buffer solution (Invitrogen, 15568025) with 25 mM EDTA (Invitrogen, 15575020), 0.5% SDS (w/v) (Fisher Scientific, BP1311), and 20 μg/mL RNase A (Sigma-Aldrich, 10109142001)) with incubation 1 hour at 37°C 1 hour followed by proteinase K (New England Biolabs, P8107S) digestion at 50°C for 3 hours. Next, we isolated DNA by mixing the sample with 10 mL of Phenol/Chloroform/Isoamyl Alcohol (Fisher Scientific, BP1752I100) with phase-lock gel (5g of Corning High Vacuum Grease (Dow Corning, 1658832) in a tube. After centrifuging at 2800xg for 10 minutes, we precipitated the DNA by adding the aqueous phase to 4 mL of 5 M ammonium acetate (Invitrogen, AM9070G) and 30 mL of ice cold 100% ethanol (Decon Labs, 2716). To pellet the DNA, we centrifuged the solution at 12,000xg for 5 minutes and performed washed with 70% ethanol twice. After leaving the DNA pellet to dry at room temperature for five minutes, we resuspended the DNA in 100 μL of 10 mM Tris-EDTA (pH 8.0), leaving the tubes to rotate at room temperature overnight.

Next, we to enable CRISPR/Cas9 cutting, we first dephosphorylated the genomic by incubation with of 5 μg of genomic DNA, 3 μL NEB rCutSmart Buffer (New England Biolabs, B6004), and 3 μL of QuickCIP enzyme (New England Biolabs, M0525S) at 37°C for 20 minutes, 80°C for 2 minutes, and finally 20°C for 15 minutes.

For CRISPR/Cas9 preparation, we used previously published crRNAs that target upstream and downstream of the *FMR1* CGG STR (**Table S7)**. We acquired the crRNAs and tracrRNA from Integrated DNA Technologies. We resuspended each RNA species to 100 μM in 10 mM Tris-EDTA (pH 7.5) (Invitrogen, AM9858). We then annealed the 2.5 μM of crRNAs and 10 μM of tracrRNAs by incubation at 95° C for 5 minutes and cooling to room temperature in a thermocycler. We use the annealed crRNA·tracrRNA pool at 10 μM to assemble Cas9 ribonucleoproteins (RNPs) assembly in 1x NEB CutSmart buffer with 62 μM HiFi Cas9 (Integrated DNA Technologies, 1081060). To allow for assembly, we left the mixture on ice for 30 minutes.

To cut the genomic DNA, we incubated the 5 μg of dephosphorylated DNA and 10 μL of the RNPs mix along with 10 mM dATP (Thermo Scientific, R0141), and 1 μL Taq polymerase (New England Biolabs, M0273) at 37°C for 60 minutes and allowed for dA-tailing of the cut blunt ends by inbuating the sample 72°C for 5 minutes. We cleaned up the DNA with the addition of 16 μL of 5 M ammonium acetate (Invitrogen, AM9070G) and 126 μL of cold 100% ethanol. To pellet the DNA, we spun the mix down at 16,000xg for 5 minutes, followed by two washes of 70%. We left the DNA pellet to dry at room temperature for 5 minutes. We resuspended the pellet in 10 mM Tris-HCl (pH 8.0) at 50°C for 1 hour and overnight rotation at 4°C. Then, we used BluePippin DNA Size Selection (Sage Science) to enrich for on target fragments with *FMR1* CGG STR using the “0.75DF 3-10 kb Marker S1” cassette definition and size range mode at 5-12 kb.

To prepare to load the DNA onto the MinION flowcell (Oxford Nanopore Technologies), we first barcoded samples for multiplexing by adding 3 μL of barcode (Oxford Nanopore Technologies, EXP-NBD104) along with 50 μL of Blunt/TA Ligase Master Mix (New England Biolabs, M0367). After an incubation of 10 minutes at room temperature, we used 50 μL of Agencourt AMPure XP beads (Beckman Coulter, A63881) to clean up the DNA and eluted into 16 μL nuclease-free water. We then ligated the barcoded DNA with Nanopore adapters with NEBNext Quick Ligation Module (New England Biolabs, E6056S) and Ligation Sequencing Kit (Oxford Nanopore Technologies, SQK-LSK109). First, we mixed 20 μL NEBNext Quick Ligation Buffer, 10 μL NEBNext Quick T4 DNA ligase, and 5 μL Nanopore Adapter Mix (Oxford Nanopore Technologies, EXP-NBD104), the Adapter Ligation Solution. We added 65 μL prepared DNA to 20 μL Adapter Ligation Solution and mixed. Then immediately, we added an additional 15 μL of the Adapter Ligation Solution and let the solution incubate for 10 minutes are room temperature. After topping off the solution to a total volume of 100 μL with nuclease-free water, we added 100 μL of 10 mM Tris-HCl (pH 8.0) and 80 μL of AMPure XP Beads for DNA clean up. After letting the sample incubate for 10 minutes, we separated the bead bound DNA with a magnet and washed the beads with 250 μL Nanopore Long Fragment Buffer two times. After air drying for 30 seconds, we eluted the DNA with 14 μL of Nanopore Elution Buffer. To prepare the DNA for loading onto the MinION flowcell, we mixed 13 μL the DNA with 37.5 μL Nanopore Sequencing Buffer and 25.5 μL loading beads. We loaded the MinION flowcell and sequenced for 48 hours.

### Cell fixation for Hi-C and ChIP-seq

We fixed cells as previously performed (*24, 85–87, 90–92, 98, 99, 103, 106–110*). We crosslinked FXS_448 iPSC-NPCs in 1% formaldehyde (ThermoScientific, 28906) in DMEM/F-12 (Thermo Fisher, 11320033) for 10 min. We quenched the fixation in 125 mM glycine for 20 min: the first five minutes at room temperature and last 15 minutes at 4 °C. We then scraped cells with a scraper (Corning, 353089) and collected the cells in PBS. We spun the cells into a pellet, froze them in liquid nitrogen, stored them at -80°C.

### Hi-C

We used the Arima Genomics Hi-C kit (Arima Genomics, A510008) to prepare the Hi-C library according to the manufacturer’s protocol as previously performed for iPSC-NPCs (*24, 103*). We lysed 2 million crosslinked cells described above and permeabilized the nuclei. We performed *in situ* enzymatic DNA digestion within nuclei. We then biotinylated ligation junctions between the digested ends. We extracted DNA and sheared it using a Covaris S220 sonicator with 140 W power, 10 % duty factor, and 200 cycles per burst for 55 seconds, resulting in fragments of an average size of ∼400 bp. We used Ampure XP beads (Beckman Coulter, A63881) to size select for sheared DNA in the range of 200 to 600 bp. We pulled down fragments with biotinylated ligation junctions using streptavidin beads from the kit. We then washed twice with wash buffer and eluted into elution buffer provided in the kit, per the kit instructions. We then eluted the DNA off the beads by resuspending the beads in 15 μL elution buffer and heating the solution at 98°C for 10 minutes. Finally, we amplified the NEBNext Ultra II DNA Library Prep Kit for Illumina (NEB, E7645S) with eight cycles of PCR according to the manufacturer’s protocol. We sequenced Hi-C libraries with 37 bp paired end reads on either an Illumina NextSeq 500 or NovaSeq 6000.

### ChIP-seq

We performed ChIP-seq on FXS_448 iPSC-NPCs as previously done(*24, 85–87, 90–92, 98, 99, 103, 106–110*) for the other iPSC-NPCs to enable harmonized analysis. For CTCF ChIP-seq, we used 10 million cells and for H3K9me3 ChIP-seq, we used 3 million cells. We lysed cells by resuspending the pellet on ice in lysis buffer (10 mM Tris-HCl, pH 8.0 (Invitrogen, 15568025), 10 mM NaCl (Invitrogen, AM9760G), 0.2% NP-40/Igepal CA-630 (Sigma-Aldrich, I8896), 1X Protease Inhibitor Cocktail (Roche, 11873580001), 1 mM phenylmethylsulfonyl fluoride (Sigma-Aldrich, 93482) for 10 min. We proceeded to homogenize the sample with 30 strokes of pestle A on ice. After pelleting the lysate at 2,500 g at 4 °C, we resuspend the pellet in 500 μL nuclear lysis buffer consisting of 50 mM Tris pH 8.0, 10 mM EDTA (Invitrogen, 15575020), 1% SDS (Fisher Scientific, BP1311), 1x Protease Inhibitor Cocktail, 1 mM phenylmethylsulfonyl fluoride and left on ice for 20 minutes. Then, we added 300 μL of IP Dilution Buffer (20 mM Tris pH 8.0, 2 mM EDTA, 150 mM NaCl, 1% Triton X-100 (Sigma-Aldrich, 93443), 0.01% SDS, 1 mM phenylmethylsulfonyl fluoride) and transferred the solution into sonication tubes. We sonicated the samples using a QSonica Q800R2 sonicator with settings of 100% amplitude, pulsed 30 seconds on and 30 seconds off, for an hour at 4 °C. Then, we spun down the sonicated lysate at 18,800xg at 4 °C.

To remove the nuclear membrane debris, we transferred the supernatant into 800 μL of pre-clearing solution. The pre-clearing solution contained 3.7 mL IP Dilution Buffer, 500 μL nuclear lysis buffer, 50 μg of rabbit IgG (Sigma-Aldrich, I8140), and 175 μL of a 1:1 ratio of Protein A (Thermo Fisher, 1591801) and Protein G (Thermo Fisher 15920010) bead slurry. We left the supernatant and preclearing solution rotating at 4°C for 2 hrs. Afterwards, we reserved 200 μL of the pre-clearing reactions as the input control. We added the remaining solution was to a new tube with 1 mL cold PBS, 20 μL Protein A, 20 μL Protein G, and either 10 μL of CTCF antibody (Millipore 07-729) or 3 μL H3K9me3 antibody (Abcam ab8898). We rotated the solutions overnight at 4°C for antibody binding.

The following day, we pelleted the beads at 2050xg for 5 minutes at 4°C and discarded the supernatant. We washed the pellet first with IP Wash Buffer 1 (20 mM Tris pH 8, 2 mM EDTA, 50 mM NaCl, 1% Triton X-100, 0.1% SDS), followed by two washes of High Salt Buffer (20 mM Tris pH 8, 2 mM EDTA, 500 mM NaCl, 1% Triton X-100, 0.01 % SDS), then another wash with IP Wash Buffer 2 (10 mM Tris pH 8, 1 mM EDTA, 0.25 M LiCl (Sigma-Aldrich, L9650), 1% NP-40/Igepal, 1% sodium deoxycholate (Sigma-Aldrich, D6750)), and finally twice more with TE buffer (10 mM Tris pH 8, 1 mM EDTA pH 8). Then, we resuspended the pellet in freshly prepared elution buffer (100 mM NaHCO_3_ (Sigma-Aldrich, S5761), 1% SDS). After spinning the beads down at 7,000 RPM, we degraded RNA with the addition of 2 μL RNase A (Sigma-Aldrich, 10109142001) and incubation at 65°C for 1 hour. Next, we reversed crosslinks by adding 3 μL proteinase K (NEB, P8107S) for incubation overnight at 65 °C. The next day, we recovered DNA with phenol:chloroform extraction and ethanol precipitation. We prepared ChIP-seq libraries using the NEBNext Ultra II DNA Library Prep Kit (NEB, E7645S) according to the manufacturer’s protocol. We used Ampure XP beads (Beckman Coulter, A63881) to size select for fragments < 1 kb in length and used 11 cycles of PCR to amplify the library. We sequenced ChIP-seq libraries with 75 bp single-end reads on an Illumina NextSeq 500.

### RNA-seq

We performed RNA-seq as previously described (*24, 103, 110*). We used the mirVana miRNA Isolation Kit (Thermo Fisher, AM1560) according to the manufacturer’s protocol to isolate RNA from iPSC-NPCs. To remove genomic DNA, we treated the isolated prep with with rDNAse I (Ambion, AM1906) according to the manufacturer’s instructions. We used RNA 6000 Nano kit (Agilent, 5067-1511) on the BioAnalyzer (Agilent) to confirm RNA samples had an RNA Integrity Number > 9. Then, we took 100 ng of RNA and prepared an RNA-seq library using the TruSeq Stranded Total RNA Library Prep Gold kit (Illumina, 20020598) per the manufacturer’s protocol. Briefly, we removed rRNA, synthesized cDNA with 0.8 U of SuperScript II RT (Thermo Fisher, 4376600) per sample, and performed A-tailing end repair. We then ligated indices from the TruSeq RNA Single Indexes Set A (Illumina, 20020492) and used Ampure XP beads (Beckman Coulter, A63881) to size select for DNA 300 bp long. We prepared the RNA-seq libraires using the TruSeq Stranded Total RNA Library Prep Gold (Illumina, 20020598) according to the manufacturer’s instructions. We sequenced RNA-seq libraries with 75 bp paired-end reads on an Illumina NextSeq 500.

### qRT-PCR

We performed qRT-PCR as previously described (*23, 24*). We isolated RNA from flash frozen iPSC pellets stored at -80°C iPSCs using the mirVana miRNA Isolation Kit (Thermo Fisher, AM1560) and treated with rDNAse I (Ambion, AM1906) according to the manufacturer’s instructions. We inputted 200 ng RNA per sample into the SuperScript III First-Strand Synthesis System for RT-PCR (Thermo Fisher, 18080-05) to convert the RNA to cDNA following the manufacturer’s protocol. For each reaction, we used 2 μL of cDNA, 400 nM forward and reverse primers (**Table S6**) in 1x Power SYBR Green PCR Master Mix (Thermo Fisher, A46109). We also created standards for each gene target ranging from 5e2 to 5e-4 pM by serial dilution. We used Applied Biosystems StepOnePlus Real-Time PCR System with the following setting for qPCR: 95°C for 10 min, followed by 40 cycles of 95°C for 15 seconds and 65°C for 45 seconds. We confirmed amplicon specificity by running a melting protocol to record a curve after each qRT-PCR reaction.

### CUT&RUN processing

We processed CUT&RUN data as previously performed to enable integrated analyses (*24*). To align the CUT&RUN data to the GRCh38 genome assembly, we used bowtie2 (version 2.2.5) with parameters “--local --very-sensitive-local --no-mixed --no-discordant --phred33 -I 10 -X 700”. We used fixmate, sort, and view “-F 4” commands from samtools (version 1.11) to remove unmapped reads and output bam files. We downsampled mapped to read depth for IgG and H3K9me3 samples to the lowest number of mapped reads with samtools view with parameters “-hbs” and set a seed of 42 for reproducibility. After index creation with samtools index, we input normalized using deeptools (version 3.3.0) bamCompare using the “–extendReads –binSize 10 –smoothLength 30 – operation log_2_” parameters to create the bigwigs.

### Hi-C processing

We mapped Hi-C reads individually to the hg38 genome assembly using bowtie2 (*111*) through Hi-C Pro (*112*) (v 2.7.7) as previously performed for the published iPSC-NPC Hi-C (*24*). We used the following parameters (--very-sensitive –L 30 –score-min L,-0.6,-0.2 –end-to-end --reorder; local parameters: --very-sensitive –L 20 –score-min L,-0.6,-0.2 –end-to-end –reorder). After filtering for unmapped reads, non-uniquely mapped reads, and PCR duplicates, we paired the resultant reads. We assembled *cis* Hi-C contact matrices at 20 kb resolution. We balanced using the Knight-Ruiz algorithm and normalized the matrices across samples by using median-of-ratios size factors conditioned on genomic distance (*24, 86*).

### ChIP-seq processing

We processed the ChIP-seq sequencing reads as previously performed for the published iPSC-NPC ChIP-seq (*24, 87, 90–92, 98, 99, 106*). We used bowtie (*113*) (v 0.12.7) to map the reads were mapped to the hg38 reference genome with parameters: “--tryhard --time --sam -S -m2”. To filter for optical and PCR duplicates, and unmapped reads, we used samtools (*114*) (v 1.11) sort, markdup -r, and view -F 4. After indexing with samtools index, to enable harmonized comparison, we downsampled across samples to match read depths using samtools function view –hbs with a seed value of 42. For bigwig generation for visualization, we used deepTools (v3.3.0) bamCoverage with default parameters for CTCF. For H3K9me3, we used bamCoverage with the additional flag, -operation subtract.

### Brain tissue CUT&RUN H3K9me3 domain calling

We used RSEG (version 0.4.9) (*115*) to define broad H3K9me3 domains from brain tissue CUT&RUN data as previously performed (*24*). We used bedtools (v2.92.2) bamtobed to convert the downsampled, filtered bam files into bed files. We ran RSEG with parameters -s 800000 -bin-size 100 -P -posterior-cutoff 0.5 -duplicates -d. We provided deadzones that we generated from RSEG deadzone using defaults parameters and a kmer size of 37.

We further refined the broad H3K9me3 domains identified by RSEG. To create a unified set of FXS brain BREACHes, we first combined RSEG calls from all N=4 FXS brain samples to create a BED file of putative BREACHes. To facilitate the identification of H3K9me3 domains present only in FXS patient brain samples and absent in the matched donor brain samples, we combined the H3K9me3 RSEG calls from all N=4 matched donor brain samples for the NL matched donor brain samples by using bedtools merge with a margin of 50 kb and then filtered for domains of at least 25 kb into a single control domain BED file. To filter the putative BREACHes based on the control domain constructed from the N=4 matched donor brain samples, we removed any overlap with control domains using bedtools subtract and merged the resultant domains within 20 kb. Finally, to retain only Megabase-scale domains, we filtered for remaining H3K9me3 domains with larger than 1 Megabase, yielding the final set of FXS brain BREACHes. To tally BREACHes in each of the N=4 FXS brain samples in a way that recognizes that at the same BREACH the extent of H3K9me3 acquisition could vary, we counted a BREACH as present if the FXS brain sample’s RSEG calls overlapped with the FXS brain BREACH by at least ten percent.

### BREACH identification from iPSC-NPC ChIP-seq H3K9me3

We used RSEG (version 0.4.9) (*115*) to define broad H3K9me3 domains from iPSC-NPC ChIP-seq data as previously performed (*24*). We used downsampled, filtered bam files into bed files using bedtools. We ran RSEG-Diff with parameters -mode 2 -s 800000 -bin-size 100 -P -posterior-cutoff 0.9995 -d.

Then for each iPSC-NPC ChIP-seq H3K9me3 samples, we merged regions within 10 kb from the initial RSEG output and filtered for regions greater than 250 kb in size. Next, we combined all regions from FXS iPSC-NPCs into a single BED file and merged regions within 50 kb to construct the putative BREACHes set. Regions from NL iPSC-NPCs were combined and merged within 10 kb to create a set of H3K9me3 domains found in NL iPSC-NPCs, our control domain regions. We then filtered regions from the putative BREACHes set that overlapped by more than 50% with control domain regions. The remaining regions were further merged within 100 kb and filtered to retain only those greater than 250 kb in size. To tally BREACHes, we counted a BREACH as present if a FXS iPSC-NPC’s RSEG calls overlapped with the FXS iPSC-NPC BREACH by at least 20%.

### MASTR-seq analysis

We performed MASTR-seq analysis as previously described (*24, 49*). We refer to the MASTR-seq manuscript for detailed analysis pipelines. We used guppy (v. 6.2.1) for base calling and barcode demultiplexing from the raw Nanopore fast5 files. We aligned read to hg38 using minimap2 (v 2.22-r1101). We proceeded with analyses on reads that aligned to the *FMR1* locus, mapped to the reverse strand, contain the known upstream sequence of the CGG tract and at least contained four executive CGGs, and did not contain nine tandem TAs. We calculated CGG STR length by counting the number of CGGs within the first and last instances of three consecutive CGGs. We then used nanopolish (v. 0.13.2) to call DNA methylation at the CpG island at the *FMR1* promoter (chrX:147911574-147912682). Briefly, we indexed the raw fast5 files the nanopolish index and called CpG methylation with call-methylation. We defined methylated CpGs as those with log_2_ likelihood > 0.1 and unmethylated as < -0.1.

### RNA-seq processing

We used kallisto (*116*) (v. 0.44.0) quant with 100 bootstraps to align RNA-seq reads to the hg38 reference. We then used tximportData(*117*) (v. 1.22.0) to convert transcript level counts to gene level counts (*24, 103, 110, 118*). We exclude genes with total counts less than 20 times the number of samples (7 iPSC-NPCs times two replicates) and performed median of ratios normalization.

### H3K9me3 ChIP-seq and CUT&RUN signal and heterogeneity quantitation

We measured H3K9me3 signal at genes genome-wide, using genes from Matched Annotation from the NCBI and EMBL-EBI (MANE) (*119*) version 1.3 for GRCh38. For each sample of H3K9me3 ChIP-seq and CUT&RUN signal, we summed the signal starting at two kilobases upstream gene’s transcription start site and ten kilobases downstream with 100 bp bins and sum all positive bins, representing signal greater than IgG negative control.

To quantify the H3K9me3 heterogeneity at a gene across the FXS patient samples (brain and iPSC-NPCs), we used the H3K9me3 signal at each gene, calculated above. We calculated to the coefficient of variation (σ/μ, CV) across the FXS patient samples within each sample type (brain and iPSC-NPCs).

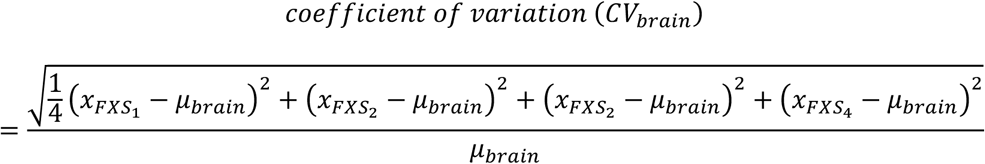

where 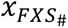 is H3K9me3 signal at each gene for each FXS patient brain sample and *μ_brain_* is the average H3K9me3 signal at each gene across the FXS patient brain samples.

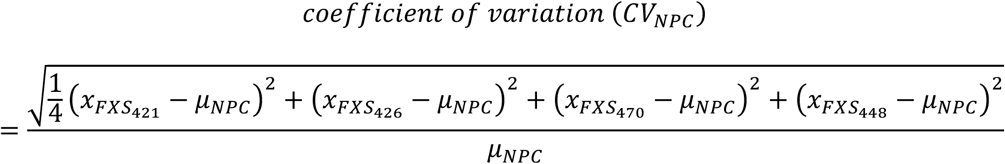

where 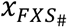 is H3K9me3 signal at each gene for FXS patient iPSC-NPCs and *μ_NPC_* is the average H3K9me3 signal at each gene across the FXS patient iPSC-NPCs.

For each of the set of samples (brain and iPSC-NPC), we converted the CV to a z-score in python using scipy (v 1.10.1). Thus, each gene had two CV z-scores: one based on the H3K9me3 heterogeneity across FXS patient brain samples (N=4) and another one based on the H3K9me3 heterogeneity across FXS patient iPSC-NPCs (N=4).

### SFARI ASD gene annotation and randomization testing

We considered genes annotated by SFARI gene scoring as “syndromic”, “category 1, high confidence”, or “category 2, strong candidate” as associated with ASD (*25*). To perform randomization testing for ASD association enrichment, we created null distribution by taking 10,000 draws without replacement with an N that matched the gene set being tested: N=20 for mosaic and invariant brain BREACH genes, N=9 for mosaic NPC BREACH genes, and N=6 for invariant NPC BREACH genes.

### Quantification of the variation in BREACH gene mRNA reduction

To quantify the heterogeneity in the extent the levels of NPC BREACH gene mRNA were reduced across the N=4 FXS patient derived iPSC-NPCs, we calculated the standard deviation for the reduction value of each NPC BREACH gene across the N=4 FXS patient derived iPSC-NPCs, each of which had two replicates. We defined the reduction value of each gene for each FXS iPSC-NPC sample to be the normalized gene mRNA count of that sample minus the average normalized mRNA count across the N=4 NL donor derived iPSC-NPCs, each of which had two replicates. If the normalized gene mRNA count of that sample was greater than the average NL normalized mRNA count, the reduction value was set to zero because that indicates the NPC BREACH gene mRNA level was not reduced in that FXS iPSC-NPC sample relative to NL iPSC-NPC samples.

### Additive models for clinical metric predictions

We wanted to evaluate the combined power of an individual gene mRNA level and FMRP level from FXS patient blood to predict clinical metrics of interest. So, we individually took each protein coding gene detectable in FXS patient blood samples from (*54*) (N= 15076 genes) and combined it with FMRP measurements (“PMBC [ng FMRP/ug total protein]”) from the same dataset. For each gene and FMRP pair, we constructed a linear regression model using the Ordinary Least Squares method with the ABC score from FXS patients using the statsmodels (v 0.14.1) in python. Thus, each gene resulted in an equation below:

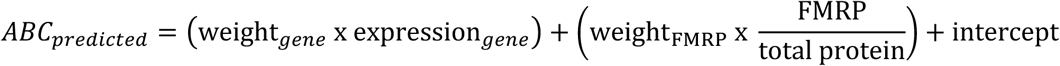

Here, we include the equations for the predictive NPC and brain FXS H3K9me3 BREACH genes and the top two predictive non-BREACH genes (**Fig. 5C&G, Fig. S3**.)

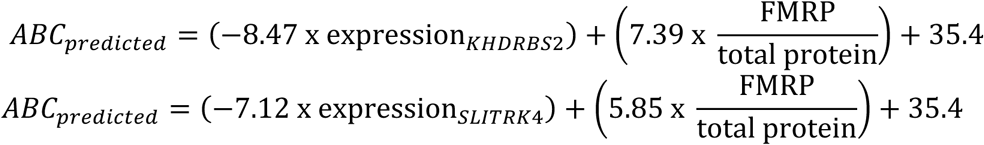

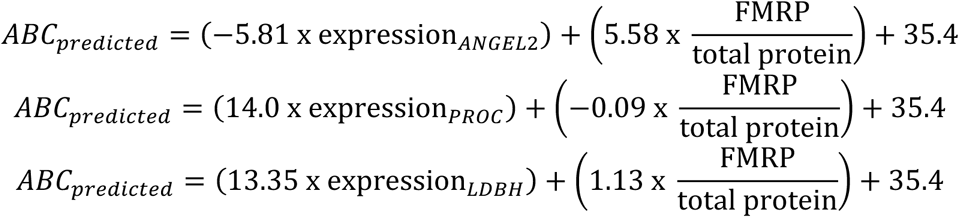

We assessed the strength of association between the gene-FMRP pair and the ABC score across FXS patients with the Pearson’s correlation coefficient (ρ) and its associated p-value using scipy (v 1.10.1). We additionally used statsmodels (v 0.14.1) to compute a q-value to control the false discovery rate by applying a Benjamini-Yekutieli correction, over a Bejamini-Hochberg correction, to be more conversative and to accommodate arbitrary dependences between the tests.

### Correlation between FMRP or gene mRNA level with clinical metric

To show correlations between blood RNA-seq gene levels and FMRP protein levels with either ABC, we use computed a Pearson’s correlation coefficient (ρ) and its associated p-value using scipy (v 1.10.1).

### BREACH genes randomization testing

We performed two sets of randomization testing. The first we tested for the average negative log10(q-value) from **Additive models for clinical metric predictions** of mosaic and invariant NPC BREACH genes compared to non-NPC BREACH genes. We sampled the non-NPC BREACH set a thousand times, each time matching the CV cutoff used for the NPC BREACH genes and the length of the gene set being tested—either the mosaic or invariant NPC BREACH gene set—representing our null distribution. We then calculated a right-sided statistic.

Second, with mosaic NPC BREACH genes that added predictive power to the linear model as defined by a q-value cutoff of 0.05, we tested the average rho for those two genes compared to non-NPC BREACH genes with a q-value greater than 0.05. To do so, we drew two non-NPC BREACH with a q-value greater than 0.05 a thousand times to create the null distribution and calculated a left-sided statistic since we were assessing the enrichment for negative correlations of the three predictive mosaic NPC BREACH genes.

### Single-nuclei RNA-seq data and ADI-R scores from individuals with ASD

We downloaded single-nucleus RNA-seq of cortical tissue from individuals with and without a diagnosis of ASD from https://autism.cells.ucsc.edu and the combined Autism Diagnostic Interview–Revised (ADI-R) results (*60*). First, we computed a combined ADI-R rank by ranking the individuals ASD across the five components the ADI-R, where higher values were ranked lower since higher values represented more severe disease. Then we summed the ranks across the five components to tally a final ranking for each individual where higher ranks represent less severe disease. Finally, to correlate gene expression from L2/3 neuronal nuclei with the combined ADI-R ranking, for everyone with ASD, we computed on a per gene basis an average log2FC of gene count in the L2/3 nuclei over average gene in all control L2/3 neuronal nuclei. We assessed correlations with a Pearson’s correlation coefficient (ρ) and its associated p-value using scipy (v 1.10.1).

## Acknowledgments

We thank Rohan Patel and all members of the Cremins lab for helpful discussion and feedback. We thank Ravi Boya for sharing qRT-PCR primers and standards and Wanfeng Gong for assistance with sequencing.

## Funding

NIH NIMH (1R01MH120269; 1DP1MH129957; JEPC)

4D Nucleome Common Fund (1U01DK127405, 1U01DA052715; JEPC)

NSF CAREER Award (CBE-1943945; JEPC)

CZI Neurodegenerative Disease Pairs Awards (2020-221479-5022; DAF2022-250430; JEPC)

NIH F30 (F30HD114405; KP)

NIH F31 Fellowship (F31NS129317; TM)

## Author contributions

Conceptualization: KP, JEPC

Methodology/Visualization: KP, TM, LZ, JHK, CS

Investigation: KP, TM, JEPC

Funding: KP, TM, JEPC

Administration: JEPC

Writing & Editing: KP, JEPC

---

**Fig. S1.**
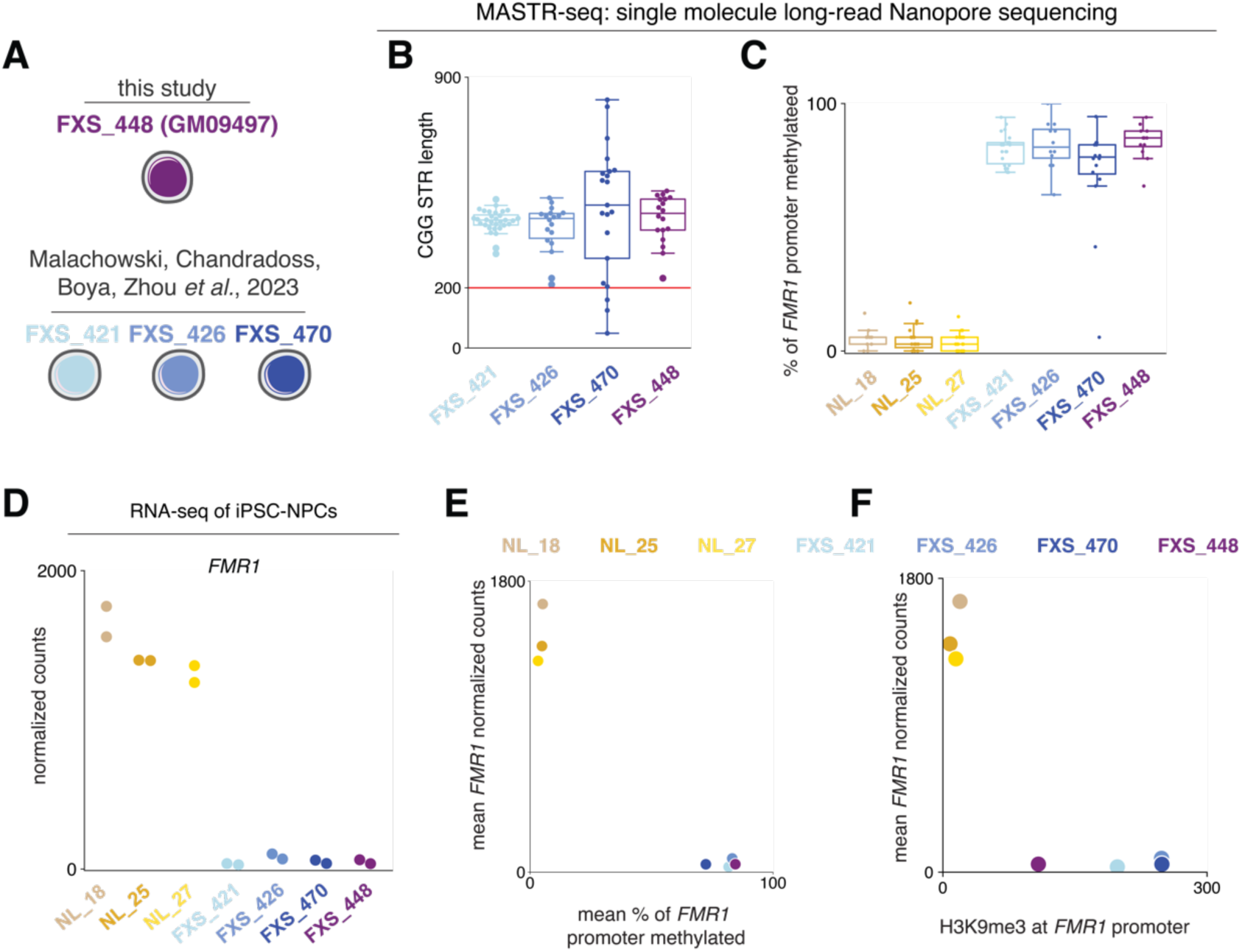
Full mutation length expansion of the CGG STR, DNA methylation at the *FMR1* promoter, and *FMR1* transcriptional silencing in FXS patient-derived iPSC-NPC lines. **(A)** Illustration of a new fourth FXS patient-derived cell line along with the previously characterized three FXS patient-derived cell lines (*24*). **(B)** CGG STR length in the four FXS iPSC lines assayed with MASTR-seq. **(C)** Percent of DNA methylation at the *FMR1* promoter across three NL iPSC lines and four FXS iPSC lines with MASTR-seq. **(D)** RNA-seq normalized counts for *FMR1* mRNA in the four FXS patient iPSC-NPCs. **(E)** CGG repeat length and DNA methylation at the *FMR1* promoter across the four FXS patient iPSCs. **(F)** CGG repeat length and H3K9me3 at the *FMR1* promoter across the four FXS patient iPSCs.

**Fig. S2.**
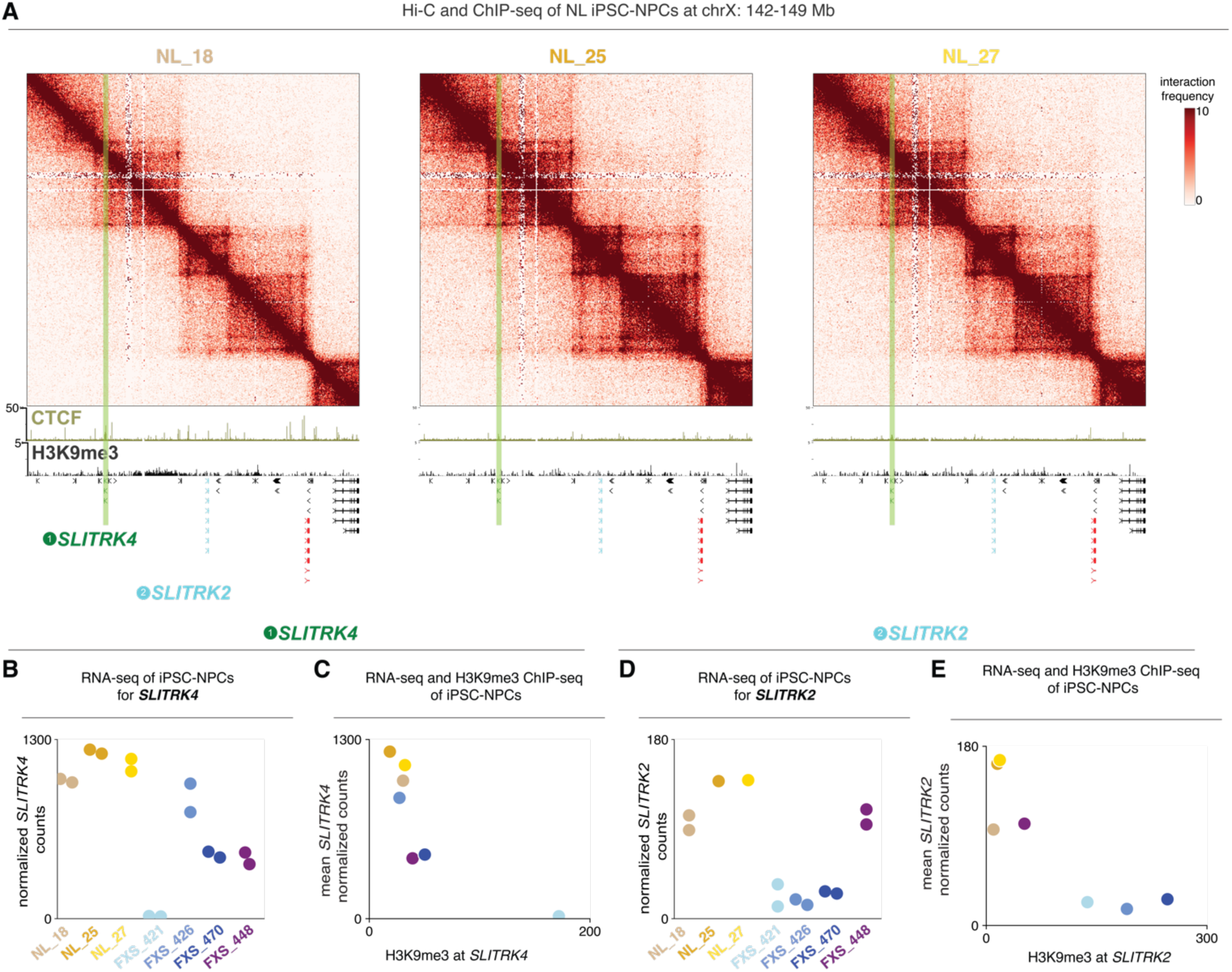
Lack of heterochromatin deposition and preserved higher-order genome folding and gene expression across NL matched donor iPSC-NPCs. **(A)** Hi-C heatmaps and CTCF and H3K9me3 ChIP-seq tracks of NL matched donor iPSC-NPCs at a 7 Mb region on the X chromosome including *SLITRK4* (green) and *SLITRK2* (blue) (*24*). **(B)** RNA-seq for *SLITRK4* mRNA in the three NL matched donor and four FXS patient iPSC-NPCs. **(C)** Relationship between H3K9me3 signal at *SLITRK4* and *SLITRK4* expression across three NL matched donor and four FXS patient iPSC-NPCs. **(D)** RNA-seq for *SLITRK2* mRNA in the three NL matched donor and four FXS patient iPSC-NPCs. **(E)** Relationship between H3K9me3 signal at *SLITRK2* and *SLITRK2* expression across three NL matched donor and four FXS patient iPSC-NPCs.

**Figure S3.**
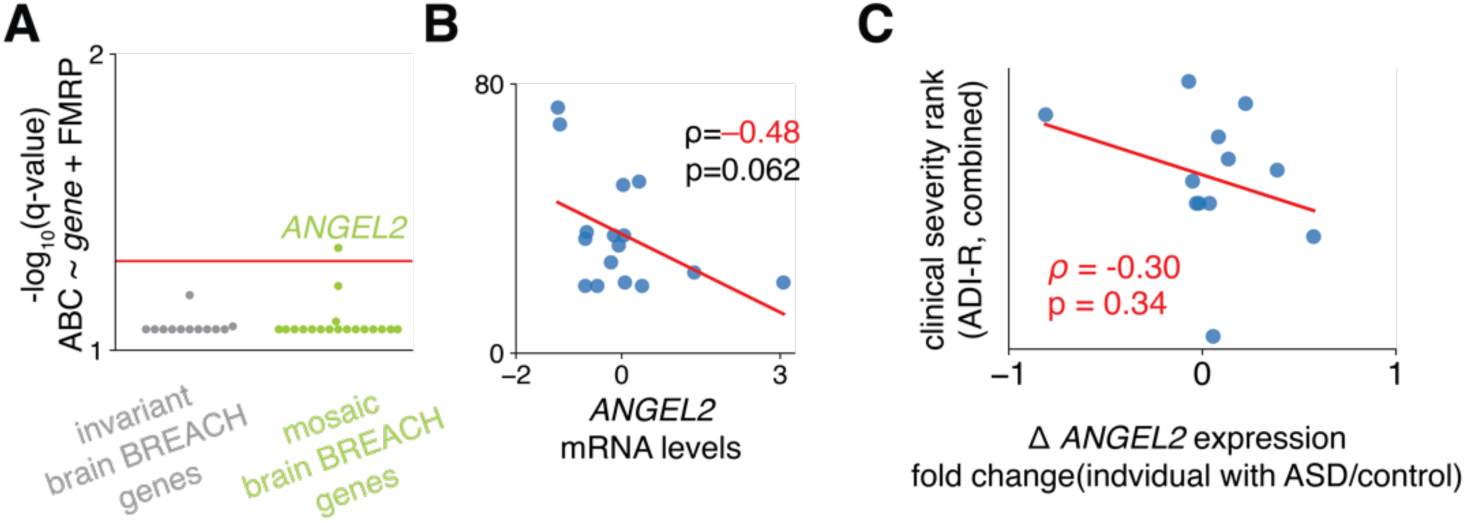
Mosaic brain BREACH gene, *ANGEL2*, is anti-correlated with FXS clinical metrics and ASD clinical metrics. **(A)** Q-values for the correlation of each gene FXS patient blood mRNA level and FMRP protein level pair with ABC score for mosaic and invariant brain BREACH genes, and genome-wide for non-BREACH genes. **(B)** Relationship between FXS patient blood mRNA levels of the most predictive mosaic brain BREACH gene, *ANGEL2,* and measure of Adaptive Behavior Composite, ABC, across FXS patients (N=16) (*54*). **(C)** Relationship between ASD patient mRNA levels of the most predictive mosaic brain BREACH genes, *ANGEL2*, and combined Autism Diagnostic Interview–Revised (ADI-R) score ranks (N=12) (*60*).

**Figure S4.**
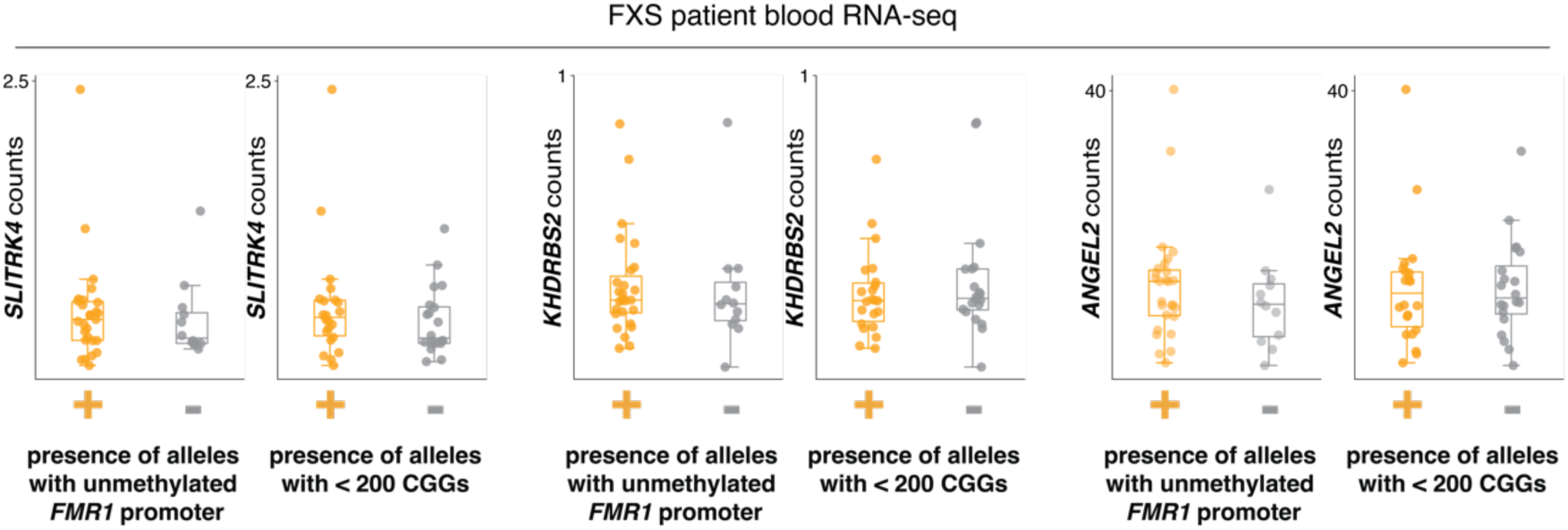
Mosaic BREACH genes correlate with FXS severity but not with established markers of clinical severity, such as size or methylation mosaicism. FXS patient blood RNA-seq (*54*) for *SLITRK4, KHDRBS2,* and *ANGEL2* stratified by methylation and size mosaicism.

## Supplementary Tables

**Table S1. Brain donor information.**

**Table S2. FXS brain BREACHes.**

**Table S3. FXS brain BREACH genes.**

**Table S4. FXS iPSC-NPC BREACHes**

**Table S5. FXS iPSC-NPC BREACH genes**

**Table S6. Transcriptome-wide in FXS patient blood evaluation of correlation between gene mRNA and FMRP levels with ABC in FXS patients**

**Table S7. qRT-PCR primers and sgRNA sequences**

